# Functional organization and natural scene responses across mouse visual cortical areas revealed with encoding manifolds

**DOI:** 10.1101/2024.10.24.620089

**Authors:** Luciano Dyballa, Greg D. Field, Michael P. Stryker, Steven W. Zucker

## Abstract

A challenge in sensory neuroscience is understanding how populations of neurons operate in concert to represent diverse stimuli. To meet this challenge, we have created “encoding manifolds” that reveal the overall responses of brain areas to diverse stimuli and organize individual neurons in stimulus–response coordinates according to their selectivity and response dynamics. Here we use encoding manifolds to compare the population-level encoding of primary visual cortex (VISp) with that of five higher visual areas (VISam, VISal, VISpm, VISlm, and VISrl), using data from the Allen Institute Visual Coding–Neuropixels dataset from the mouse. We show that the topology of the encoding manifold for VISp and for higher visual areas is continuous, with smooth coordinates along which stimulus selectivity and response dynamics are organized with layer and cell-type specificity. Surprisingly, the manifolds revealed novel relationships between how natural scenes are encoded relative to static gratings—a relationship conserved across visual areas. Namely, neurons preferring natural scenes preferred either low or high spatial frequency gratings, but not intermediate ones. Analyzing responses by cortical layer reveals a preference for gratings concentrated in layer 6, whereas preferences for natural scenes tended to be higher in layers 2/3 and 4. The results demonstrate how machine learning approaches can be used to organize and visualize the structure of sensory coding, thereby revealing novel relationships within and across brain areas and sensory stimuli.

**Significance Statement:** Manifolds have become commonplace for analyzing and visualizing neural responses. However, prior work has focused on building manifolds that organize diverse stimuli in neural response coordinates. Here, we demonstrate the utility of an alternative approach: building manifolds to represent neurons in stimulus/response coordinates, which we term ‘encoding manifolds.’ This approach has several advantages, such as being able to directly visualize and compare how different brain areas encode diverse stimulus ensembles. This approach reveals novel relationships between layer-specific responses and the encoding of natural versus artificial stimuli.

## Introduction

Understanding how populations of neurons are organized to represent external stimuli is a core goal of sensory neuroscience. This is made nontrivial by the fact that both artificial and natural stimuli are typically high dimensional, as are the patterns of spiking activity that these stimuli produce across populations of neurons. Such complexity poses a challenge to visualizing, let alone understanding, how complex stimuli are represented by neural population activity. To address this, we have recently developed *encoding manifolds* to organize neurons according to their responses to an ensemble of visual stimuli [24]; these manifolds allow one to visualize how neural populations encode ensembles of stimuli. In the encoding manifold, each point is a neuron, and nearby neurons respond similarly in time to members of the stimulus ensemble. The encoding manifold differs from more standard applications of manifolds in neuroscience, in which each point on the manifold is a stimulus (or a point along a trajectory) embedded in neural response coordinates (see, e.g., [15] for a review). This distinction is relevant as the latter approach emphasizes reading out the stimulus (or movement trajectory) from the population response (i.e., decoding), while the former organizes neurons according to their functional properties (i.e., encoding), which can drive hypotheses about cell types and their anatomical and functional connectivity in a circuit.

One feature of encoding manifolds exploited in prior work [24] is they allow for comparing the topology of sensory encoding at different stages of sensory processing. When comparing the encoding manifolds between retina and primary visual cortex (VISp) in mouse to an ensemble of artificial stimuli (drifting gratings) and more naturalistic stimuli (flows), they were strikingly different: the manifold inferred from retinal responses consisted of largely separate clusters of neurons that corresponded to known retinal ganglion cell types. This result served to lend credence to the encoding manifold as a useful approach for revealing underlying biological structure because the stimuli used to generate the manifold were distinct from those used to classify the retinal ganglion cell types. The VISp manifold derived from responses to the same stimuli was much more continuous than the retinal manifold, with neurons effectively carpeting stimulus/response space. Like the retinal manifold, the structure of the VISp manifold was quite informative about the physiological organization of VISp. While no information about laminar position or broad-spiking versus narrow-spiking neurons (putative excitatory vs. inhibitory neurons, respectively) was used in its construction, these properties were not randomly located across the manifold; rather, functional cell types were distributed in a complementary fashion across different layers and in different regions.

This prior work raised several important questions: to what extent does the structure of the VISp encoding manifold depend on the stimulus set? To what extent are the encoding manifolds of higher visual areas similar to that observed in VISp? Finally, can the encoding manifold reveal aspects of how natural scenes are encoded by visual cortex? Fortunately, these three questions could be answered by utilizing the Allen Institute’s large dataset of visual cortical responses of VISp and higher cortical visual areas to a variety of stimuli [59, 19, 20]. The stimulus set includes stationary gratings of a wide range of orientations, spatial frequencies, and spatial phases, a set of a similar number of stationary natural images, and an ensemble of drifting gratings across a range of temporal frequencies and orientations (Fig. 1a,b). Using these data, we have constructed encoding manifolds for each of the visual areas using only the responses to static and drifting gratings. The resulting VISp encoding manifold was similar to that in our earlier work despite substantive differences in the stimulus sets. Namely, both the present and previous manifolds were continuous, with foci for different cortical layers and broad-spiking versus narrow-spiking cell types. The higher visual areas also exhibited similar manifold structures. One major similarity among the manifolds generated from each visual area was a preference for static gratings over natural scenes in layer 6, a layer that projects back to sub-cortical structures and elsewhere [53]; neurons in all other layers, across all visual areas, tended to prefer natural scenes.

**Fig. 1:**
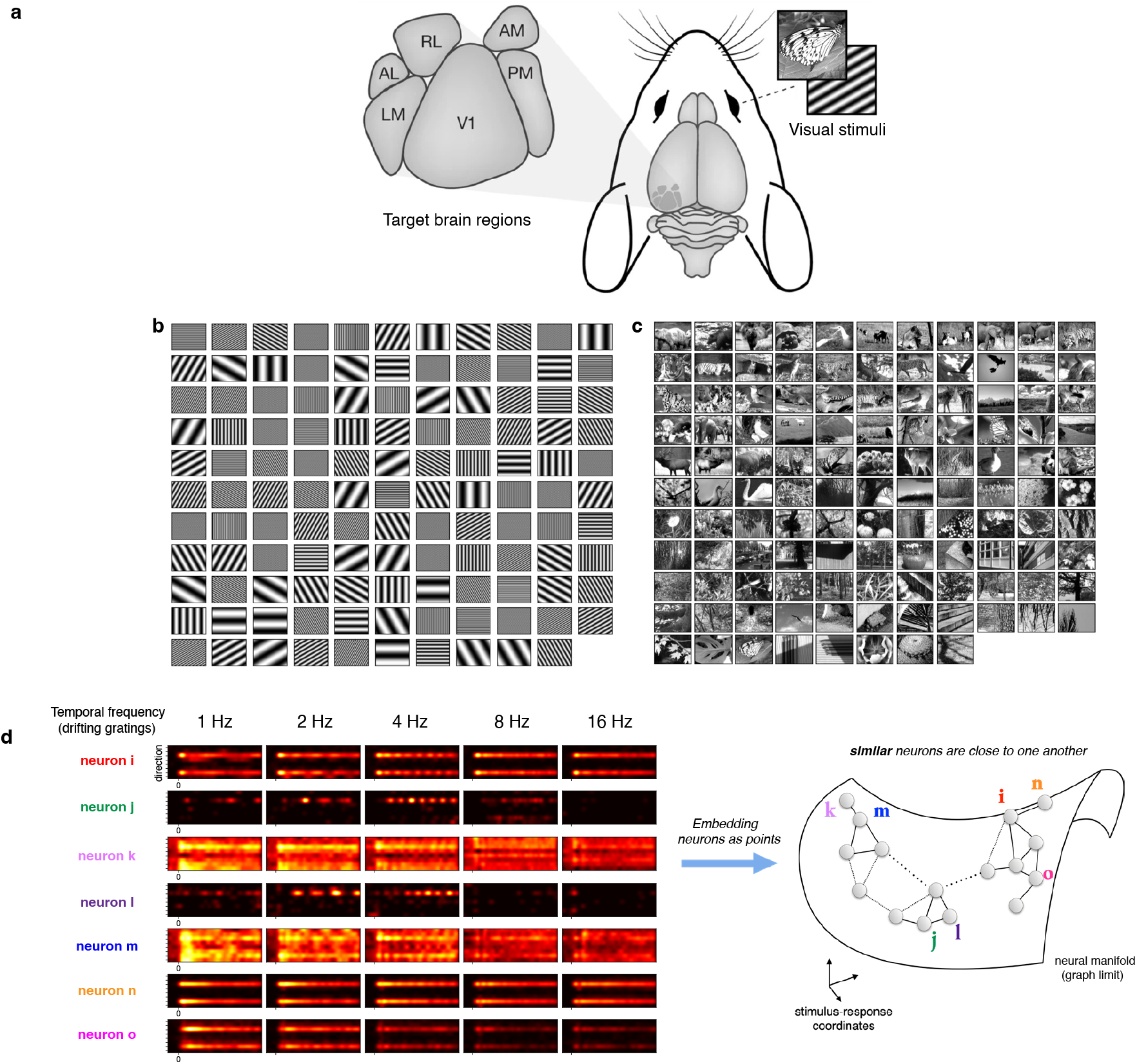
Building encoding manifolds from visual cortical responses to artificial and naturalistic stimuli. **a**, The locations of the visual cortical areas recorded at the Allen Institute while presenting drifting gratings, flashed static gratings, and flashed natural scenes. **b–c**, The ensemble of static gratings (**b**) and natural scenes (**c**) used in the Allen dataset. **d**, PSTHs of responses to an ensemble of drifting gratings that vary in temporal frequency (columns) for several neurons (rows). The encoding manifold organizes neurons so that those nearby on the manifold respond similarly in time to similar stimuli. Neurons *i* and *n* exhibit similar activity patterns, and are close, while neurons *i* and *j* are different. This is illustrated by the underlying data graph (see Methods) superimposed on the manifold. Panel **a** modified from [20]; panel **d** after [24].

Further analysis of the natural scene responses using the encoding manifold revealed a particularly interesting geometry. The resulting manifolds had an axis that ran from neurons with strong to weak orientation selectivity. However, those neurons that responded better to natural images relative to gratings formed a band in the manifold roughly orthogonal to this axis in V1 (referred to as VISp in the Allen Institute data and below), which suggests that there exist natural-scene preferring neurons over the full range of selectivity for stimulus orientation. Furthermore, our encoding manifolds revealed that natural-scene preferring neurons also preferred either high or low spatial frequencies rather than intermediate ones, which were preferred by the neurons driven best by gratings. This structure was also present in each of the higher visual areas, although the axes differed from VISp. These results further validate the use of encoding manifolds for assaying the structure and topology of sensory coding across distinct stimulus ensembles and stages of sensory processing, and reveal novel aspects of the global organization of neurons across the visual system.

## Results

We began by determining to what extent the structure of the encoding manifold of mouse primary visual cortex (VISp) generalizes from our prior visual stimulus set to that utilized by the Allen Institute [4]. In previous work, we analyzed responses to drifting gratings and naturalistic ‘flow’ stimuli, composed of points and line segments moving semi-coherently [23, 24]. However, the previous data were limited to VISp and did not thoroughly examine how the structure of the encoding manifold depended on the temporal frequency composition of the stimulus. The Allen Institute dataset has several features that allow a useful comparison (Fig. 1). First, it consists of measurements not only from VISp but also from five higher cortical visual areas in mouse: VISam, VISal, VISpm, VISlm, VISrl (Fig. 1a). Second, the stimulus set contains drifting gratings presented at five temporal frequencies and eight directions. Third, it contains static gratings presented at multiple spatial frequencies (five), orientations (six), and phases (four) (Fig. 1b). Finally, it contains static images of natural scenes (Fig. 1c). Thus, we used the Allen Institute dataset to generate, analyze, and compare encoding manifolds along the mouse cortical visual hierarchy.

### The encoding manifold reveals novel relationships between stimulus selectivity and neuronal classes

Encoding manifolds were calculated from the Allen Institute data using the responses to both drifting and static gratings. In brief, the encoding manifold arises from a machine learning algorithm that learns to place individual neurons in a relatively low-dimensional space according to their similarity in stimulus selectivity and response dynamics: neurons nearby on the manifold exhibit similar temporal responses to similar stimuli (Fig. 1d) [24]. Responses to the natural scene stimuli were not included in the calculations of the encoding manifolds because the embedding algorithm requires stimuli for which neuronal responses do not depend on receptive field location (see Methods).

Consistent with prior results [24], the encoding manifold of VISp was relatively smooth and continuous, with neurons uniformly carpeting stimulus/response space (Fig. 2a). Note that this topology is quite distinct from the manifold produced by this analysis when applied to retinal responses [24]. Representative peri-stimulus time histograms (PSTHs) for 5 temporal frequencies are shown for several neighborhoods of neurons; differences in selectivity for stimulus direction and in the temporal responses to drifting gratings across the manifold are hinted at by these PSTHs.

**Fig. 2:**
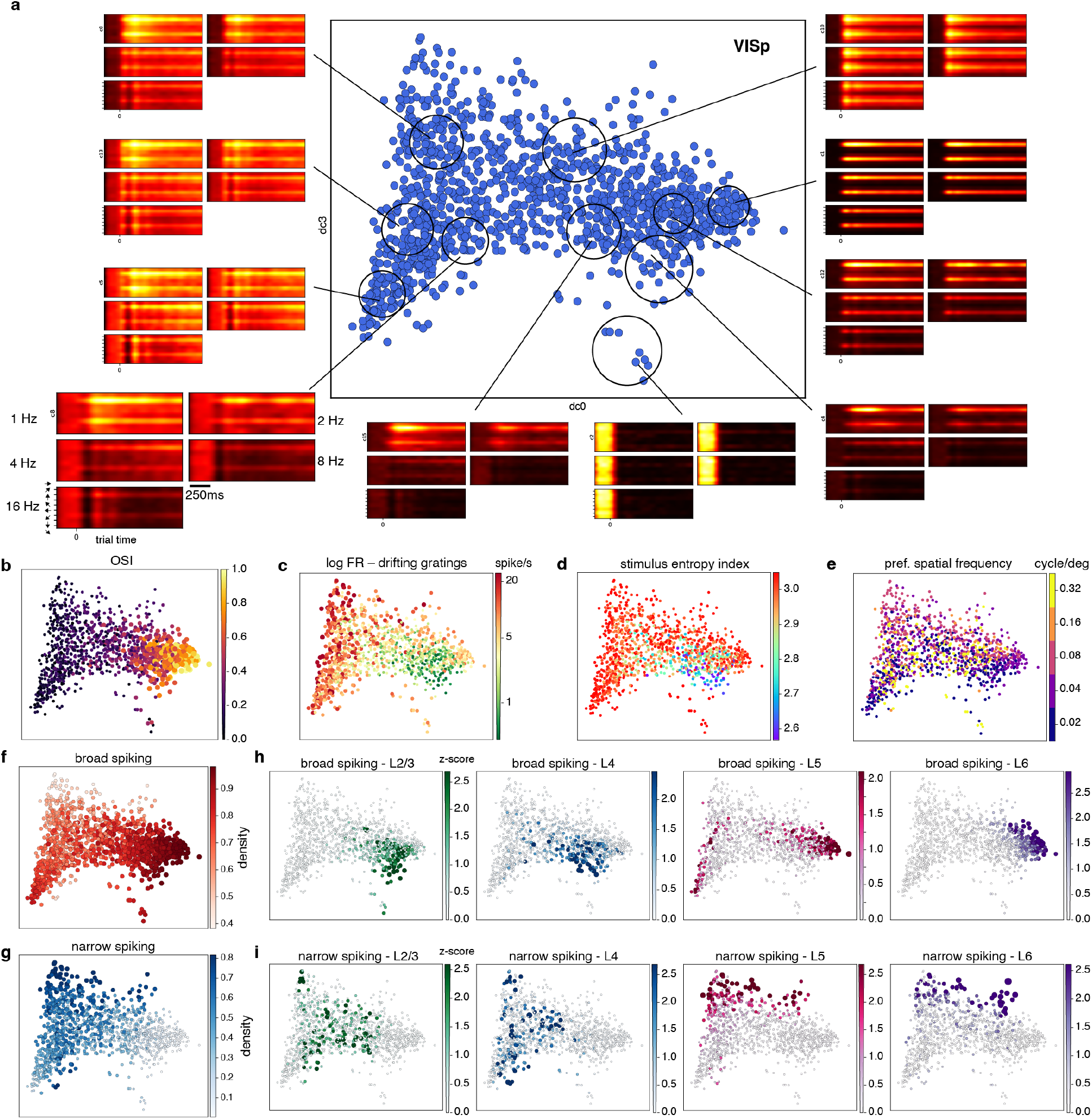
The encoding manifold for VISp (V1) computed from responses to gratings. **a**, A projection onto the first (dc0) and fourth (dc3) diffusion coordinates of the encoding manifold (see Methods) shows a smooth transition in ensemble PSTHs, from neurons sharply tuned for orientation (right side) to those broadly tuned (left side), indicating a continuous topology. The mean PSTHs for 10 different groups of neurons indicated on the manifold are arranged by temporal frequency from left to right then top to bottom at 1, 2, 4, 8, and 15 Hz. The PSTHs have been aligned by the preferred orientation to de-emphasize the specific orientation to which each neuron is tuned. **b–d**, Encoding manifold in **a** reproduced, with points colored according to orientation selectivity index (OSI) (**b**), firing rate (**c**), stimulus entropy (**d**), and preferred spatial frequency (**e**). Note the OSI values vary smoothly and continuously across the manifold, but are progressively less organized for stimulus entropy and preferred spatial frequency. **f–g**, Encoding manifold from **a** with points colored by the local density of broad-spiking neurons (**f**) or narrow-spiking neurons (**g**): note the distributions are complementary. **h–i**, Encoding manifolds reproduced showing the locations of broad-spiking and narrow-spiking neurons by cortical layers 2/3, 4, 5, and 6.

To more clearly visualize how response properties are organized across the manifold, neurons (points) can be colored according to their preferred stimuli and/or response features. Clear gradients or geometries emerged along particular coordinates. For example, a gradient from left to right in the orientation selectivity index (OSI) of each cell was revealed by the manifold (Fig. 2b), as well as one for firing rate increasing from lower right to upper left (Fig. 2c). However, not all stimulus features were so well organized. For example, entropy, a measure of the range of stimuli for which a neuron gives robust responses, varied only slightly across the manifold (Fig. 2d), while neurons were not obviously organized by their spatial frequency preference (Fig. 2e; a point we will return to later). These observations underline an advantage of the encoding manifold: for a given firing rate, or orientation selectivity, there is a range of spatial frequencies and stimulus preferences. Without the global organization of the manifold, such co-variation would be difficult to discern.

Broadly, the organization of stimulus features across the manifold had a similar topology to that from our previous study that used a distinct stimulus ensemble [23, 24]. This suggests that the VISp encoding manifold is not strongly dependent on the chosen stimulus set, provided that the set is sufficiently diverse. Of course, we cannot rule out that other topological features (i.e., distinct clusters) might emerge with yet-to-be tested stimuli. The continuous topology across the manifold is markedly different from that of the retina [24], in which morphologically and functionally distinct retinal ganglion cell types form distinct clusters on the manifold. This difference suggests that cell type diversity in visual cortex serves a distinct purpose than in the retina, as discussed previously [24].

The organization of features that were not used in the calculation of the manifold is more interesting. Neurons were classified on the basis of their electrical waveforms: broad-spiking or narrow-spiking (typically treated as putative excitatory and putative inhibitory neurons, respectively, although we note that some broad-spiking cortical neurons are known to be inhibitory and express somatostatin [44]). The broad-spiking neurons were concentrated on the right in this view of the manifold, with a secondary concentration salient at the lower left, and were present at a lower concentration throughout the middle of the manifold (Fig. 2f). In contrast, the narrow-spiking neurons were concentrated in the upper left corner and were also present at a lower concentration throughout the middle of the manifold; they were less densely represented in the regions of the manifold in which the broad-spiking neurons were concentrated (Fig. 2g). Thus, the encoding manifold recapitulates the observation that narrow-spiking neurons tend to be less selective for stimulus orientation and to fire at higher rates than broad-spiking neurons in mouse V1 [54, 34]. Again, this confirms that the encoding manifold organizes neurons by their physiological properties.

The encoding manifold for V1 also organized neurons according to cortical layers even though layer information was not used to generate the manifold (Fig. 2h,i). In layers 2/3, broad-spiking neurons were concentrated in the lower right, as were those in layer 4 though less tightly. In layer 6, broad-spiking neurons were much more tightly concentrated in the upper right. It is known that layer 6 broad-spiking neurons tend to have low firing rates [68], which is confirmed by the manifold (compare Fig 2h layer 6 to panel c). In layer 5, broad-spiking neurons exhibited two foci of density: one together with those of layer 6, but more tightly focused, and another in the salient concentration at the lower left. This bimodal distribution reveals that layer 5 contains (at least) two populations of broad-spiking neurons with quite different response properties [35] (see Discussion).

In comparison, the distributions of narrow-spiking neurons exhibited distinct patterns from the broad-spiking neurons across layers. In layers 5 and 6, narrow spiking neurons were most concentrated along the top of the manifold, whereas those in layers 2/3 and 4 were concentrated lower down except for a cluster at the very top left corner. While the layers differed from one another in the distributions of broad-spiking and narrow-spiking neurons, within each layer these two classes of neurons were largely complementary. Again, we emphasize that these features of the data were not utilized in generating the encoding manifold, and thus their recovery serves as validation that the manifold captures meaningful functional organization. At the same time, the finer within-layer structure demonstrates the ability of the manifold approach to reveal novel relationships between stimulus selectivity, response dynamics, and neuronal classes. As a control analysis, we tested whether the neurons’ receptive field position or size (see Methods) could trivially explain some of the organization found on the manifolds, but found that these properties were not well organized on any of the manifolds studied (Fig. S1).

Importantly, these results on broad-spiking and narrow-spiking neurons across layers recapitulate several observations from our previous use of encoding manifolds to understand mouse retina and VISp [24]. However, they add to that work by demonstrating that the approach is robust when the experiments use a different stimulus ensemble, different neuronal recording technologies, and are performed by different laboratories.

### Encoding manifolds from higher cortical visual areas are continuous like VISp

Do higher cortical visual areas in the mouse exhibit similar encoding manifolds with similar topology as VISp? Are these higher visual areas also continuous in their topology, and how are features such as spatial frequency, orientation, and firing rate organized across each of these neural populations? The Allen Institute data set allowed us to answer these questions for five higher visual areas: VISam, VISal, VISpm, VISlm, and VISrl.

We constructed the encoding manifolds for all five of the higher visual areas using stimuli and methods identical to those described for VISp. We found that the manifolds for all higher visual areas were continuous and shared many properties with VISp (Fig. 3). When the orientation selectivity index (OSI) and firing rate were plotted as functions on the manifolds, they varied smoothly (Fig. 3, left and center columns). High firing rates were organized with low OSI, and conversely. Finally, like VISp, spatial frequency preference did not appear well-organized for any of the higher visual areas (Fig. 3 right column), although temporal frequencies were smoothly organized along one of the diffusion coordinates (Fig. S2). Note that the apparent shape of each manifold in the two-dimensional plots depends on the specific projection used for visualization. We chose coordinates to emphasize the smooth organization of neuronal properties over each manifold (see Methods).

**Fig. 3:**
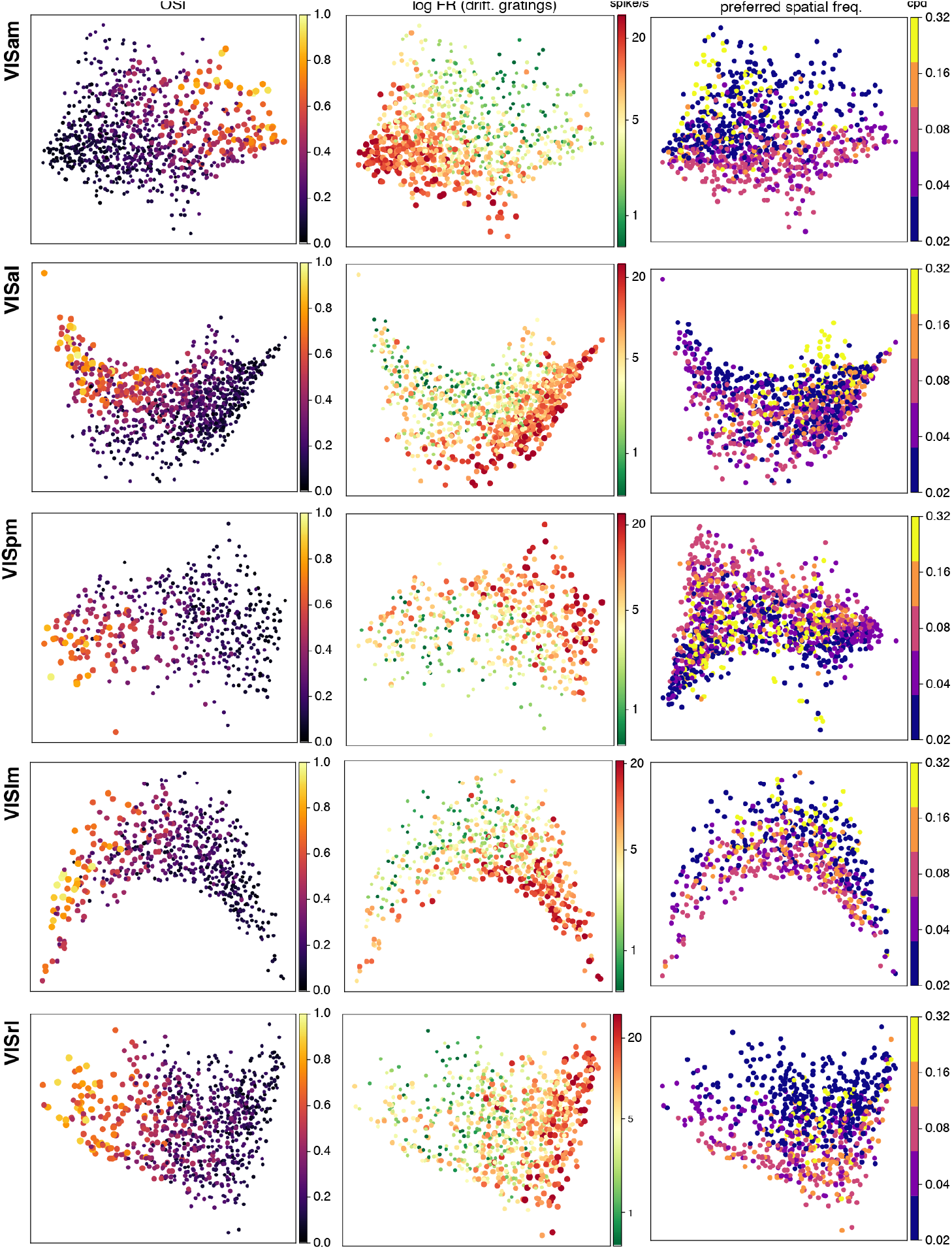
Encoding manifolds for five higher visual areas (HVAs) computed from responses to gratings. Rows indicate the HVAs and columns indicate different features: OSI and firing rate to drifting gratings (left and middle columns), and preferred spatial frequency (third column). Note the similarity of organization of these properties across areas and with VISp (Fig. 2).

We also analyzed how broad-spiking and narrow-spiking neurons were distributed across the manifolds for the higher visual areas. Similar to VISp, both classes of neurons exhibited layer-specific localization on their respective manifolds (see Figs. S4–S8), however the details of these arrangements varied somewhat across higher visual areas. One common trend was that broad-spiking neurons in layer 6 exhibited high OSIs across cortical areas. Another common trend was that broad-spiking and narrow-spiking neurons were located on complementary portions of the encoding manifold in each higher visual area and in each layer. A variable property was the bimodality of broad-spiking neurons in layer 5 observed in VISp: it varied across higher visual areas (Fig. S9).

The conclusion from this initial analysis of the encoding manifolds from the higher visual areas is that they exhibited many features in common with VISp in terms of manifold topology and geometry. Topologically, all the manifolds were continuous, distinct from the encoding manifold of retinal ganglion cells, which was more discrete (shown previously [24]). Their geometries were also similar to that of VISp, with OSI and firing rates being clearly organized across the manifolds, while preferred spatial frequencies were less clearly organized. We note these statements pertain to generating the manifolds from the responses to black-and-white gratings of various spatial and temporal frequencies; probing other stimulus features (e.g., color) could reveal greater differences across cortical visual areas (see Discussion).

### Encoding manifolds organize natural scene responses relative to gratings responses

We next examined a novel feature of the Allen Institute dataset relative to previous applications of the encoding manifold: the inclusion of responses to natural scenes. This feature provided an opportunity to examine the extent to which neural responses to natural scenes exhibited any organization on these encoding manifolds, and whether this organization varied across cortical visual areas. While we did not utilize the responses to natural scenes in generating the encoding manifolds, we could display how neurons were organized on the grating-derived manifolds according to their responses to natural scenes [32]. Given the (perhaps) unbounded variability of natural scenes [64, 45], it is difficult to know how to appropriately arrange the responses, so we began by simply plotting the mean firing rate to natural scenes across the manifold (Fig. 4a). Surprisingly, the geometric organization of the natural scene responses, like that of the grating responses, varied rather smoothly across the manifold, approximately from right to left (Fig. 4b). Comparing the kinetics of the natural scene to grating responses in individual neurons reveals similar dynamics (Fig. 4c,d), but some neurons exhibited larger responses to natural scenes (Fig. 4c), while others responded more strongly to gratings (Fig. 4d).

**Fig. 4:**
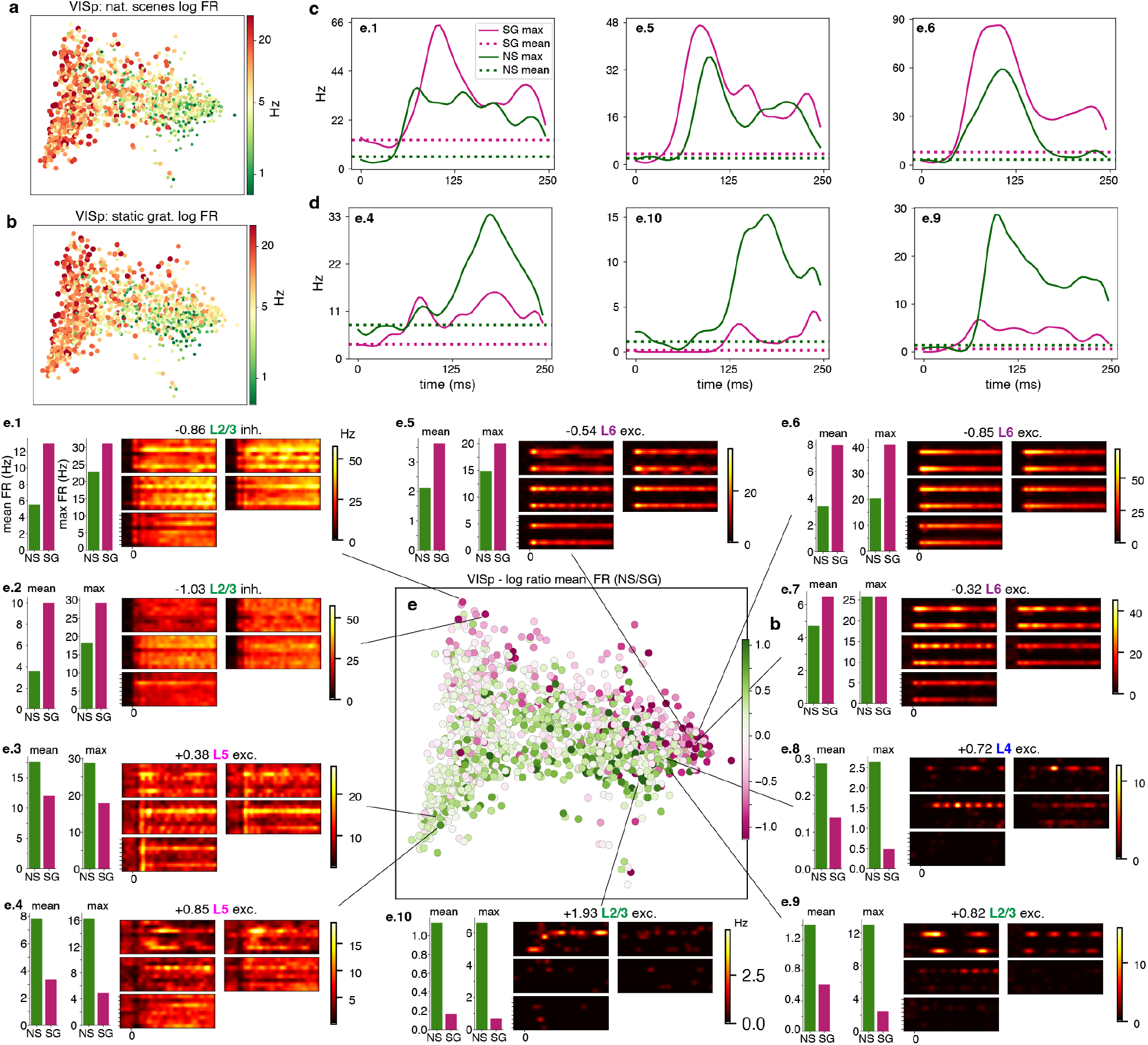
The encoding manifold inferred from gratings organizes the ratio of natural scene to static grating responses. **a–b**, Neurons on the encoding manifold from Fig. 2 colored by log mean firing rate to all natural scenes (**a**) and log mean firing rate to all static gratings (**b**). **c–d**, Firing rates (kernel-smoothed, see Methods) over at least 39 trials as a function of time for three representative neurons that respond better to static gratings than to natural scenes (**c**), or respond better to natural scenes than to static gratings (**d**); the labels shown within each plot correspond to the neuron identifiers used in panels e.1–e.10. **e**, Encoding manifold with points (neurons) colored by the ratio of their mean response to gratings versus natural scenes: note a gradient from top to bottom of neurons preferring gratings to those preferring natural scenes. **e.1–e.10**, PSTHs as in Fig. 2 and bar plots of firing rates to gratings and natural scenes are shown for representative neurons whose positions are indicated on the manifold. Bar plots show the result is robust when using either ‘mean’ (using all stimulus variations) or ‘max’ (using the preferred stimulus variation) firing rates. Notice that the top-to-bottom organization of natural scene preference is approximately orthogonal to that of firing rate and orientation selectivity (cf. Fig. 2).

It should be noted that the static grating stimuli can be said to tile the (static) stimulus space for gratings, encompassing the relevant range of orientations and spatial frequencies so as to stimulate all grating-responsive neurons well, regardless of the location of their receptive fields on the stimulus screen. Therefore, the best response measured with these stimuli for each neuron is likely to be the near-optimal grating response. However, the natural scenes are a small selection from all possible natural scenes and, since they are flashed (not drifted), the responses to them presumably depend strongly on the location of each neuron’s receptive field in the visual field.

Nevertheless, the comparable co-variation in the geometries of organization of natural scenes to gratings in the manifolds (Fig. 4a,b) inspired us to consider the ratio of each neuron’s responses to the two types of stimuli. This comparison produced a striking result (Fig. 4e): the relative responses to natural scenes were organized along an axis orthogonal to the gradient of orientation selectivity on the manifold (Fig. 2b). Specifically, the average response of each neuron to the 118 natural scenes (Fig. 1c) was compared to its average response to all the static grating stimuli (Fig. 1b). Note that the two types of stimuli were presented identically, in separate blocks that were interleaved between the repetitions of each block. Neurons that preferred natural scenes (green) were distributed along one side of the manifold while those that preferred static gratings were more prevalent on the other side (magenta). An alternative analysis compared the maximum response to any individual stimulus from among the sets of 120 static gratings and 118 natural scenes; this is potentially a noisier measure of selectivity than using the average responses to the entire stimulus set, but it produced the same result: when the manifold was colored by the ratio of each neuron’s response to the best natural scene versus its response to the best static grating, the picture was very similar (Fig. S11). This manifold’s surprising organization of natural scenes (relative to gratings) is not a trivial consequence of differences in overall firing rates: the distributions of average firing rates to static gratings and to natural scenes were similar (Fig. 4a,b).

Despite the marked difference in the two stimulus sets, a relationship between grating preferences and natural scene preferences was revealed by the encoding manifold. We note that it has been notoriously difficult to find any consistent characterization of natural scene responses in visual cortex [25, 58, 59, 10, 73], especially when considering average responses [51]. Yet the encoding manifold has found an organization whereby across the entire range of firing rates there is a range of preferences for gratings versus natural scenes.

We next checked to see if the encoding manifolds for the higher visual areas exhibited a similar geometry, organizing the ratio of natural responses to grating responses. Remarkably, every higher cortical visual area exhibited a similar organization (Fig. 5 and Fig. S10 third column). In fact, a prediction from this observation is that if a manifold were generated by co-embedding all the neurons from all the visual areas, neurons from individual visual areas should be indistinguishably mixed (as opposed to clustered by visual area). We tested this prediction and, as expected, they were highly mixed, while main properties like OSI, FR, and ratios remaining largely well organized (see Fig. S12).

**Fig. 5:**
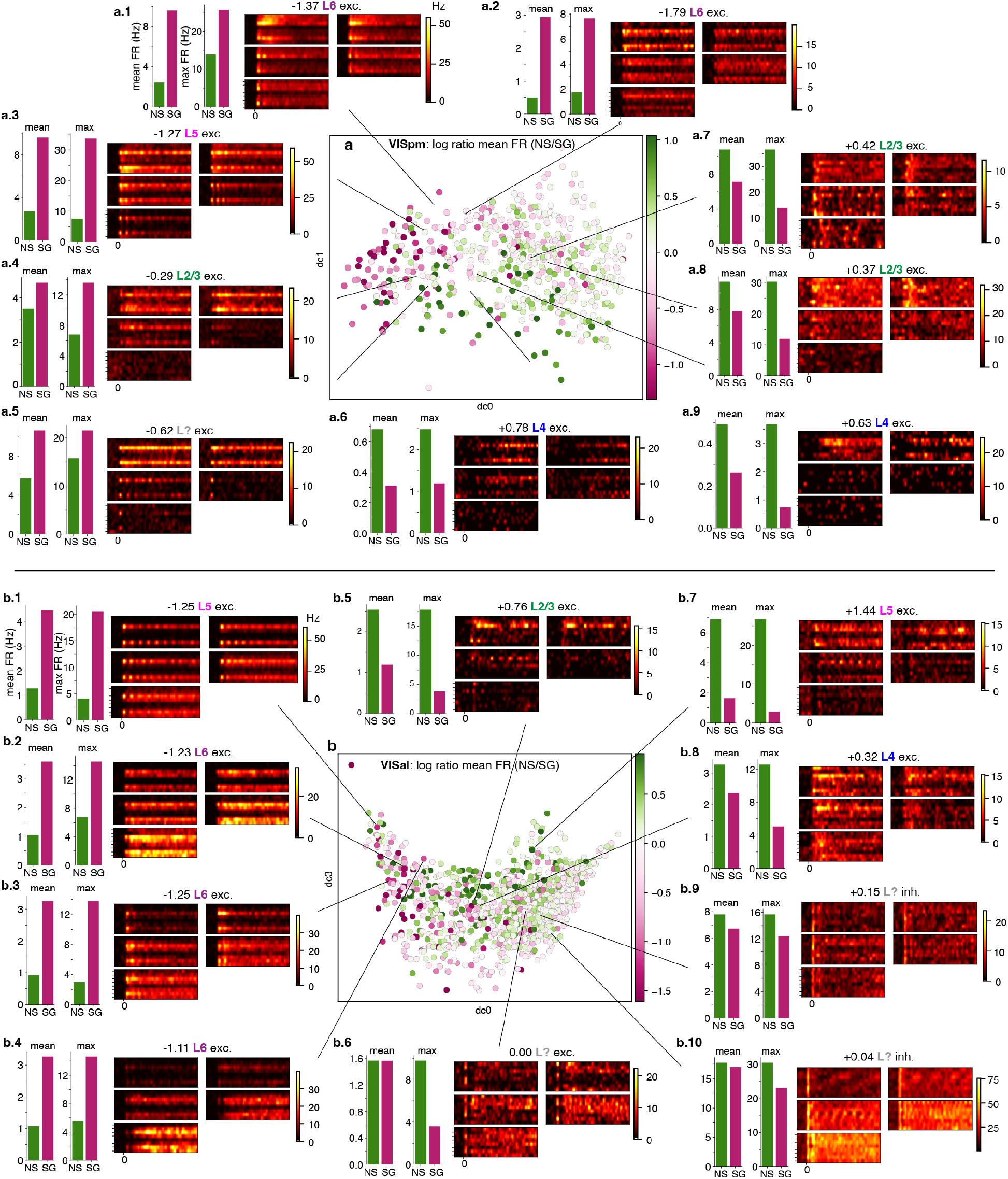
Encoding manifolds reveal preference for natural scenes vs. static gratings in VISpm (top) and VISal (bottom). **a**, The encoding manifold for VISpm colored to indicate the ratio of responses to natural scenes versus gratings. Note the gradient varying from top left (preference for gratings, in purple) to bottom right (preference for natural scenes, in green). Details of responses of nine representative neurons (**a.1–a.9**) shown as in Fig. 4e. **b**, The encoding manifold for VISal labeled by ratio of responses to natural scenes versus gratings. Note the gradient varying from left (preference for gratings) to right (preference for natural scenes). Details of responses of ten representative neurons (**b.1–b.10**) shown as in Fig. 4e.

The consistency of these results across visual cortical areas suggests something important about this organization. Furthermore, it had to be revealed by some property of the responses to only gratings because the natural scenes were not used in generating the encoding manifolds. We next sought to find this property of the grating responses that organized the preferences for natural scenes (relative to gratings).

### Global relationship between spatial frequency tuning and natural scene preferences

To uncover the signifier in the grating responses that organized the natural scene preferences, a reasonable place to start is with the selectivity to spatial frequencies. We began with VISp: as noted above, preferred spatial frequency appeared mixed over the VISp manifold (Fig. 2d). However, we examined the density of neural responses by frequency band (cf. Fig. S2) and a surprising organization was revealed upon closer inspection: there were complementary densities of neurons preferring either low or high spatial frequencies versus neurons preferring intermediate spatial frequencies (Fig. 6a). Furthermore, the spatial distributions of these preferences over the manifold varied almost identically to the distributions of neurons that preferred natural scenes versus gratings (Fig. 6b,c and Fig. S13). Thus, the population of natural-scene-preferring neurons was distributed on the manifold in a similar manner to neurons preferring either high or low spatial frequencies.

**Fig. 6:**
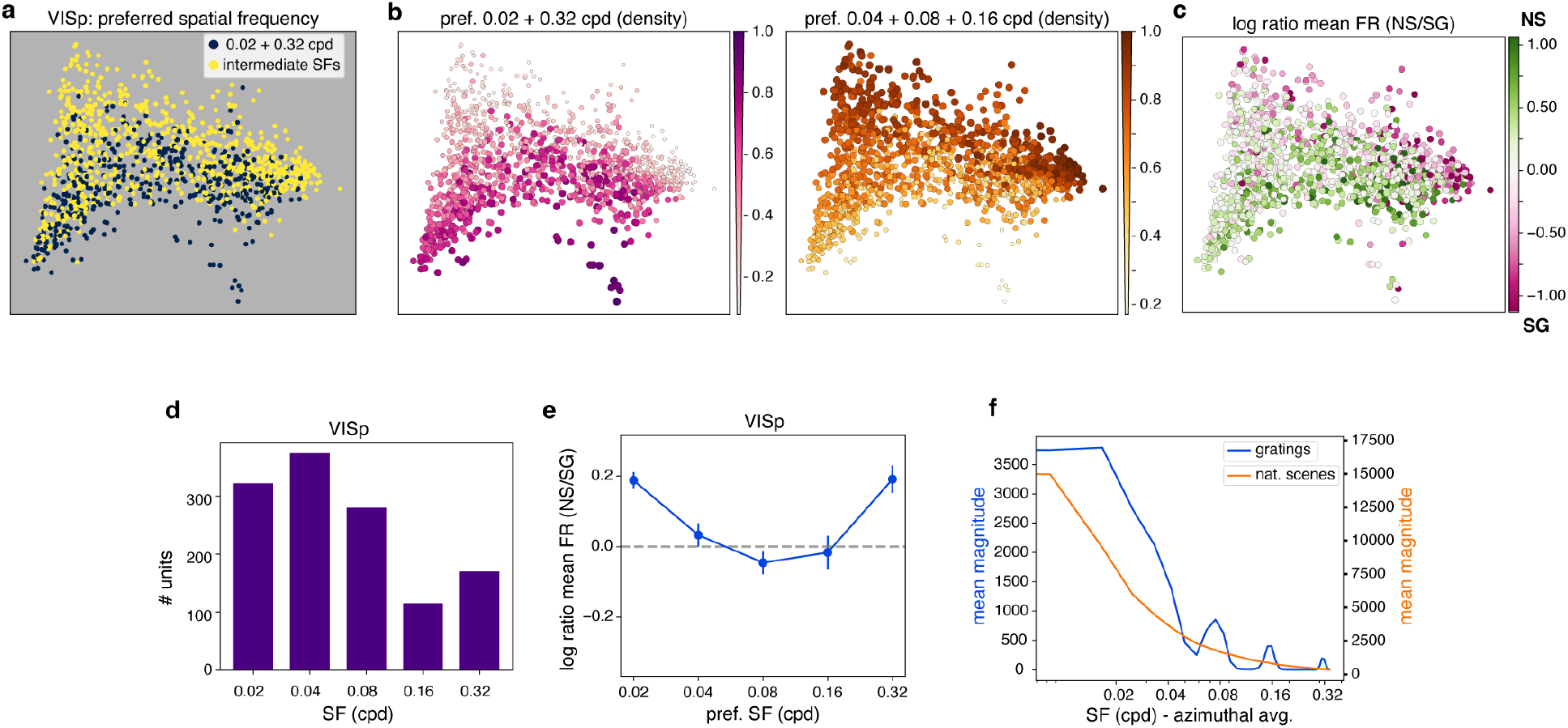
Comparing preference for spatial frequency and natural scenes versus gratings in VISp. **a**, Encoding manifold for VISp colored by spatial frequency preference: neurons are colored according to their preferred spatial frequency for static grating stimuli, grouping 0.02+0.32 (extreme frequencies, in yellow) vs. 0.04+0.08+0.16 (intermediate frequencies, in blue). **b**, Spatial frequency preferences summarized by their local density over the manifold (see Methods); note that extreme frequencies (left) and intermediate frequencies (right) are largely complementary. **c**, The log ratios from Fig. 4e, shown here for comparison with the distributions in **b**. Corresponding plots for higher visual areas are shown in Fig. S13. **d**, Distribution of number of units preferring a given spatial frequency (SF) in VISp. **e**, Comparison across neurons binned by their preferred spatial frequency (*x*-axis) of the mean (*±* s.e.m.) of the log ratios between mean firing rates to natural scenes vs. static gratings. Neurons that prefer low or high SFs tend to respond more strongly to natural scenes, while those that prefer intermediate SFs respond more strongly to gratings. **f**, Mean azimuthally averaged radial Fourier power spectra of the flashed gratings and natural scenes used in the experiments. Spectra were averaged across all stimulus instances within each stimulus class and are plotted as a function of spatial frequency on a logarithmic axis.

To quantify this relationship between grating preferences and natural scene preferences, we first binned neurons in each visual area by their spatial frequency preferences (Fig. 6d). Consistent with previous work, there was a preponderance of neurons that preferred lower spatial frequencies [26, 47]. We then calculated the log ratio of the mean response across natural scenes to the mean response across static gratings for each neuron as a function of that neuron’s preferred spatial frequency (in response to drifting gratings); the result was a “U”-shaped curve, showing that neurons in the group preferring either low or high spatial frequencies tended to have the largest responses to natural scenes, while those preferring intermediate frequencies tended to respond more vigorously to gratings (Fig. 6e). Importantly, the average radial spatial-frequency302 spectra of the grating and natural-scene stimulus ensembles were highly similar (Pearson correlation, r= 0.9037; cosine similarity, 0.9043; Fig. 6f), indicating that the higher responses to natural scenes at the extreme SFs are not trivially explained by higher energy in those stimuli at 0.02 and 0.32 cpd. Rather, neurons in mouse V1 tuned to high or low spatial frequencies tend to respond better to natural scenes than gratings, while neurons tuned to intermediate spatial frequencies are more responsive to gratings. We return to this point and its potential significance in the Discussion.

Next, we determined to what extent this relationship observed in VISp between spatial frequency preferences and natural scene preferences held for the higher visual cortical areas. Indeed, each visual area exhibited a “U”-shaped relationship between spatial frequency preference and natural scene preference (Fig. 7). Furthermore, as with VISp, this relationship was reflected in the encoding manifolds distributing neurons that preferred low+high spatial frequencies on one side, and neurons that preferred intermediate spatial frequencies on the other side of the manifold (Fig. S13). As a complementary visualization, Fig. S18 shows the mean firing rates to natural scenes and static gratings separately as a function of preferred spatial frequency for each visual area.

**Fig. 7:**
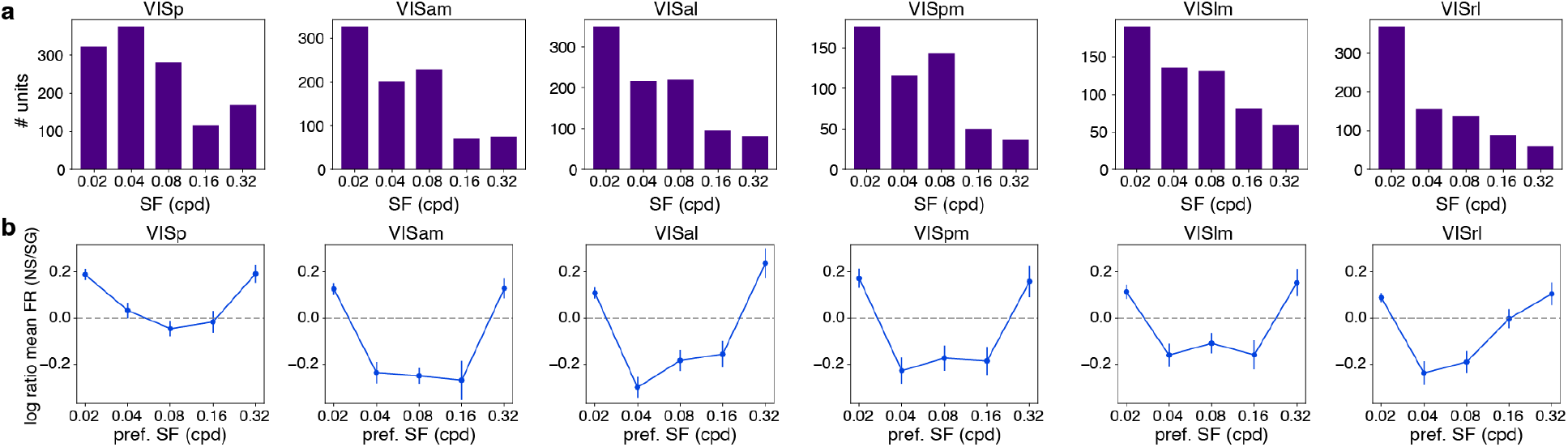
Preference for spatial frequency and for natural scenes versus gratings across visual areas. **a**, Distribution of number of units preferring a given spatial frequency (SF) in each visual area studied. **b**, Mean *±* s.e.m. log ratio between mean firing rate to natural scenes (NS) vs. static gratings (SG) of the neuronal population in each area as a function of their preferred SF. Neurons that prefer low or high SFs tend to respond more strongly to natural scenes (positive ratios), while those that prefer intermediate SFs respond more strongly to gratings (negative ratios).

Finally, to rule out the possibility that the observed relationship was merely a consequence of the gradient in firing rates to static gratings, we a permutation test. Shuffling natural-scene firing rates across neurons destroyed both the original manifold organization and the characteristic U-shaped relationship between preferred spatial frequency and the NS/SG ratio (see Fig. S14). The observed U-statistic (see Methods) exceeded that expected under the permutation null distribution in five of the six visual areas studied, indicating that the observed U-shaped relationship is unlikely to arise from chance associations between preferred spatial frequency and natural-scene responses.

### Natural scene preference differs by layer

We next checked for layer-specific preferences for natural scenes versus static gratings across visual cortical areas (Fig. 8), because some of the laminar distributions observed showed clear overlap with grating– or natural-scene-preferring regions on the manifolds of several visual cortical areas (Fig. 8a–d). To quantify this, we generated histograms of the log ratio of mean firing rates to natural scenes vs. gratings for neurons in each layer and visual area (layers 2 and 6 shown in Fig. 8e). Histograms shifted toward positive values indicate a preference for natural scenes, while a shift toward negative values indicates a preference for static gratings. As with our previous comparisons between natural scene and static grating responses, the data were analyzed in two ways. First, the mean response across all displayed natural scenes was compared to the mean response across all static gratings (‘mean analysis’). Second, the mean response to the single natural scene that generated the largest response was compared to the mean response to the single static grating that generated the largest response (‘max analysis’).

**Fig. 8:**
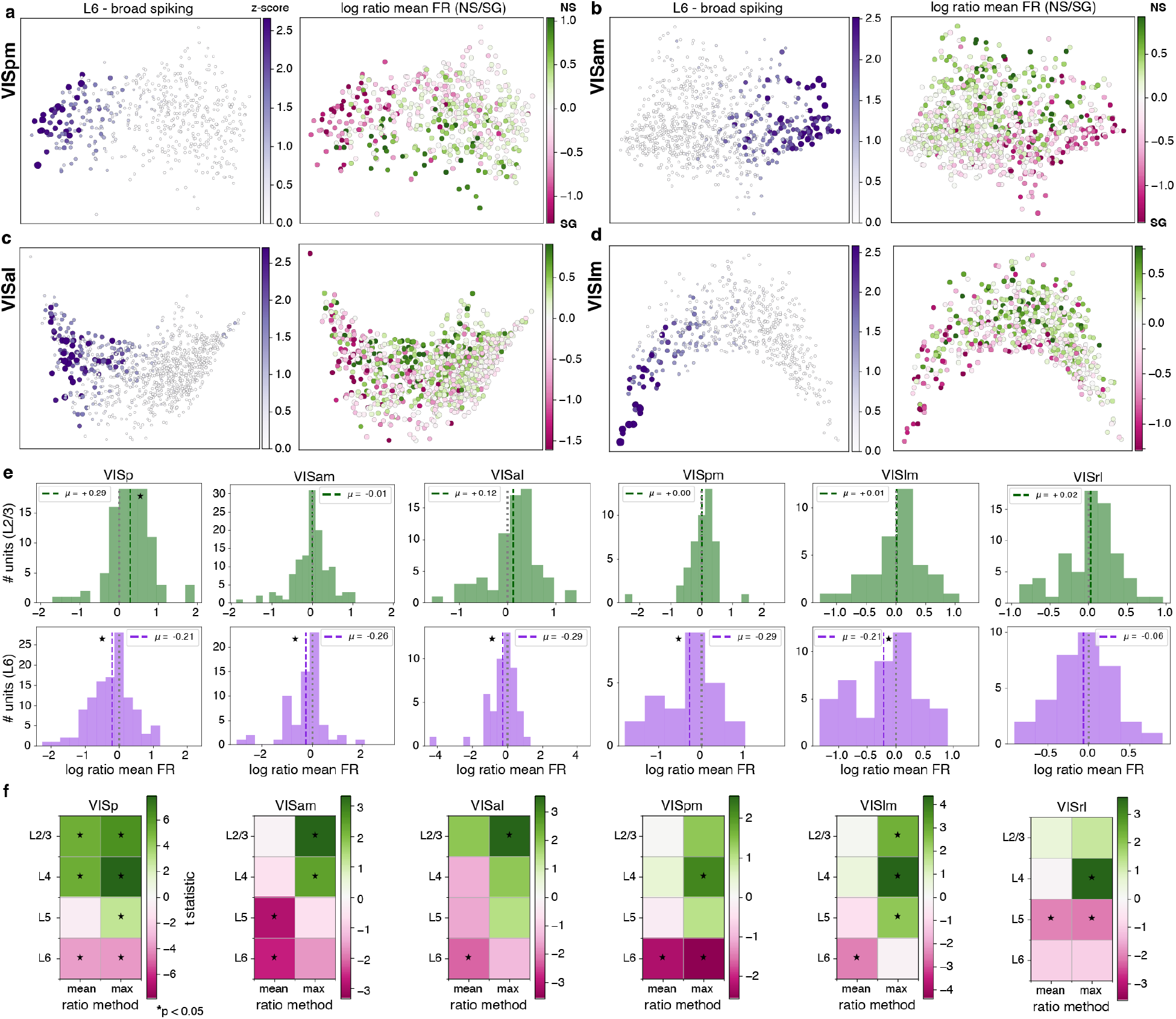
Preference for natural scenes versus gratings across cortical areas and layers. **a–d**, Local density of broad-spiking neurons belonging to cortical layer 6 (left) and preference for gratings vs. natural scenes (right) for neurons in the VISpm, VISam, VISal, and VISlm manifolds. Note the overlap between the regions where layer 6 neurons concentrate and the regions with higher preference for gratings. **e**, Histograms of the distribution of log ratios computed from the mean firing rates to natural scenes versus static gratings for all neurons in layer 2/3 (**top row**) and in layer 6 (**bottom row**); each column shows a different visual cortical area; *µ* indicates the mean. In layer 2/3, the VISp histogram is shifted toward positive values, indicating a preference for natural scenes; all other areas have histograms centered approximately around zero, indicating no layer-specific preference for natural scenes compared with gratings. By contrast, neurons in layer 6 show a significant preference for gratings in all areas (negative shift) except for VISrl. One-sample *t*-tests were performed to determine if each distribution differed significantly from a normal distribution (same variance) centered at 0. Significance (*p <* 0.05) is indicated by *★*. **f**, Summary of *t*-test statistics comparing the log ratio between natural scenes (positive, green) and static gratings (negative, magenta) responses across cortical layers and visual areas. Values computed using either the mean (left column) or maximum firing rates (right) are shown. Across visual areas, preference for natural scenes is most consistently observed in superficial layers, whereas preference for gratings tends to be stronger in deeper layers.

The ‘mean analysis’ revealed a preference among layer 6 neurons for static gratings over natural scenes in each cortical area (Fig. 8f, left columns: magenta (green) indicates grating (natural scene) preference). This preference was less pronounced under the ‘max analysis’ (Fig. 8f, right columns). In VISp, the ‘mean analysis’ revealed that layers 2/3 and 4 tend to have stronger responses to natural scenes. This was not the case in the higher cortical areas for layers 2/3 and 4. However, the ‘max analysis’ did indicate a rather consistent preference for natural scenes over static gratings for layers 2/3 and 4. We emphasize that these preferences are represented in the average performance across the population of neurons, but that within each cortical layer and cortical area, there was substantial heterogeneity (as revealed by the histograms in Fig. 8e).

## Discussion

The encoding manifold organizes large populations of neurons according to their responses to an ensemble of stimuli, thereby incorporating both circuit and stimulus properties. Responses to gratings collected by the Allen Institute confirmed the continuous topology of the encoding manifold for VISp that we found earlier using a different stimulus ensemble [24]. This lends validation to the approach as different visual stimuli and different neural recording technologies were used and the experiments were performed in a different laboratory.

The manifold analysis also revealed patterns beyond what was already known. First, higher visual areas in mouse shared a similar continuous topology with VISp. Throughout this work, we use the term “continuous” descriptively to refer to the absence of clearly separated clusters and the smooth variation of physiological properties across the manifold. A rigorous topological characterization of manifold continuity remains an interesting problem for future study [37]. Second, the functional organization in response to gratings was remarkably conserved across the visual hierarchy. Third, the manifolds revealed a surprising and consistent relationship between natural scene (NS) vs. static grating (SG) preferences. Fourth, the manifolds revealed a relationship between NS preference and spatial frequency (SF) tuning: neurons preferring NSs were concentrated at low or high SFs, while intermediate SF neurons favored SGs. Fifth, these relationships were found across all visual areas and most prominently in layer 6, which showed a consistent bias toward gratings. Sixth, the manifolds revealed a bimodal distribution of layer 5 broad-spiking neurons (one lobe overlapping with layer 6, another distinct). Finally, the complementarity of broad– vs. narrow-spiking neurons was conserved across layers and higher visual areas. These findings readily provide testable hypotheses—for example, that the two layer 5 populations correspond to distinct sub-cortical or cortico-cortical projections with different roles.

We discuss further two aspects of the Allen Institute data revealed by the encoding manifold.

### The functional organization of laminar preferences differ across visual areas

Differences in grating selectivity between visual areas have been observed previously [26, 47], and have been interpreted as a hint toward dorsal/ventral stream models [70, 63, 43]. The encoding manifold approach allowed us to refine the population analysis to individual layers, where further differences emerged, consistent with previous observations of laminar specialization in visual cortical responses [41]. For example, in VISp, the greatest concentrations of highly orientation-selective, broad-spiking neurons were found in layers 5 and 6 (Fig. 2b,g). The same was true for narrow-spiking neurons in layers 4 and 2/3, even though they occupied regions practically disjoint from those of layers 5 and 6. In contrast, although in VISal the broad-spiking neurons of layer 2/3 were in the regions of the manifold with high orientation selectivity, those in layer 4 were not (Fig. S5c,f). The same was true to a lesser extent in VISpm (Fig. S6c,f). This is perhaps surprising considering that the dominant feedforward projection from superficial layers of VISp goes to layer 4 of higher cortical visual areas [59]; one would expect the recipient layer 4 populations not to lose selectivity.

The deeper layers are also interesting, since they involve both cortical and sub-cortical projections [68]. Projections to the dLGN may sharpen receptive fields in the mouse [11] and cat [52, 60, 6]), while the cortico-cortical projection could be involved in generating endstopping and other surround effects [21]. Layer 5 also projects to the pulvinar [39], a projection that could be relevant to coordinating natural scene responses [75, 9].

Layers 5 and 6 showed different organizations for broad– and narrow-spiking neurons, depending on the area. In VISp (Fig. 2g), layer 5 exhibited two distinct concentrations of broad-spiking neurons, one of which overlapped almost completely with layer 6 in the region of highest orientation selectivity. We do not know which if either of these broad-spiking clusters may contain somatostatin-expressing interneurons. Similarly, broad-spiking layer 5 neurons in the VISpm manifold were organized in two clusters, although the overlap of the cluster with the highly selective layer 6 neurons was less complete (Fig. S6). On the other hand, in VISal, layer 5 had practically no overlap with layer 6, even though the latter remained highly selective (Fig. S5). This difference between VISpm and VISal is striking, and consistent with previous reports of functional and anatomical differences between VISal– and VISpm-related visual pathways (cf. [36]). The distributions of laminar density for VISam, VISlm, and VISrl are shown respectively in Figs. S4, S7, and S8. Layer 5 in all three cases had a bimodal distribution on the respective manifolds, with one lobe remaining consistent with the density of broad-spiking neurons in layer 6. Fig. S9 summarizes all of the above comparisons between layers 5 and 6.

In summary, building encoding manifolds from responses to diverse stimulus sets and then painting the locations of broad– and narrow-spiking neurons from different layers can provide insights and generate novel hypotheses about the common themes and distinct specialization in the functional organization of visual cortical areas. Expanding these approaches to population recordings from primate visual cortical areas may also provide important insights to the conserved versus divergent features of visual processing across species.

### A novel relationship between spatial frequency tuning and natural scene preferences

Natural scenes have been among the most difficult stimuli to analyze, and the encoding manifold revealed a unique invariant at the population level. Although the manifolds computed in this study were based solely on gratings, a consistent relationship emerged between the natural scene preferences of high– and low-spatial frequency preferring neurons compared to neurons preferring intermediate frequencies. Specifically, neurons tuned to low or high spatial frequencies tended to generate larger responses to natural scenes than gratings, while neurons tuned to intermediate spatial frequencies tended to generate larger responses to gratings (Fig. 7b). This highlights the usefulness of the encoding manifold as an organizational and visualization tool: although in principle these relationships could have been found by directly plotting log ratios vs. preferred SF, inspection of how physiological properties and feature selectivity are distributed across the manifold provides a data-driven route to discovering such relationships and understanding how they are structured across cortical areas and layers (e.g., [8]). As we show in Figs. S15 and S16, a naive analysis based on examining histograms based on standard properties like OSI and firing rate alone would not have revealed these relationships.

How might this curious relationship be explained (cf. [73, 10])? One possibility is that it has to do with object content, or at least a proxy for it. Examining the statistics of image content is instructive (Fig. 9a–d and Fig. S17). The organization of neurons preferring low and high spatial frequencies relative to intermediate frequencies (Fig. 9e,f) closely parallels the distribution of neurons preferring natural scenes versus gratings (Fig. 9g), providing a visual interpretation of the U-shaped relationship shown in Fig. 7b. Low frequencies signal “where” an object might be, while high frequencies signal “what” it might be made of. Such an identification of spatial frequency content with scene layout has long been proposed [65, 56]. However, it is unlikely that simply extracting filtered features is the complete story (e.g., see [71]).

**Fig. 9:**
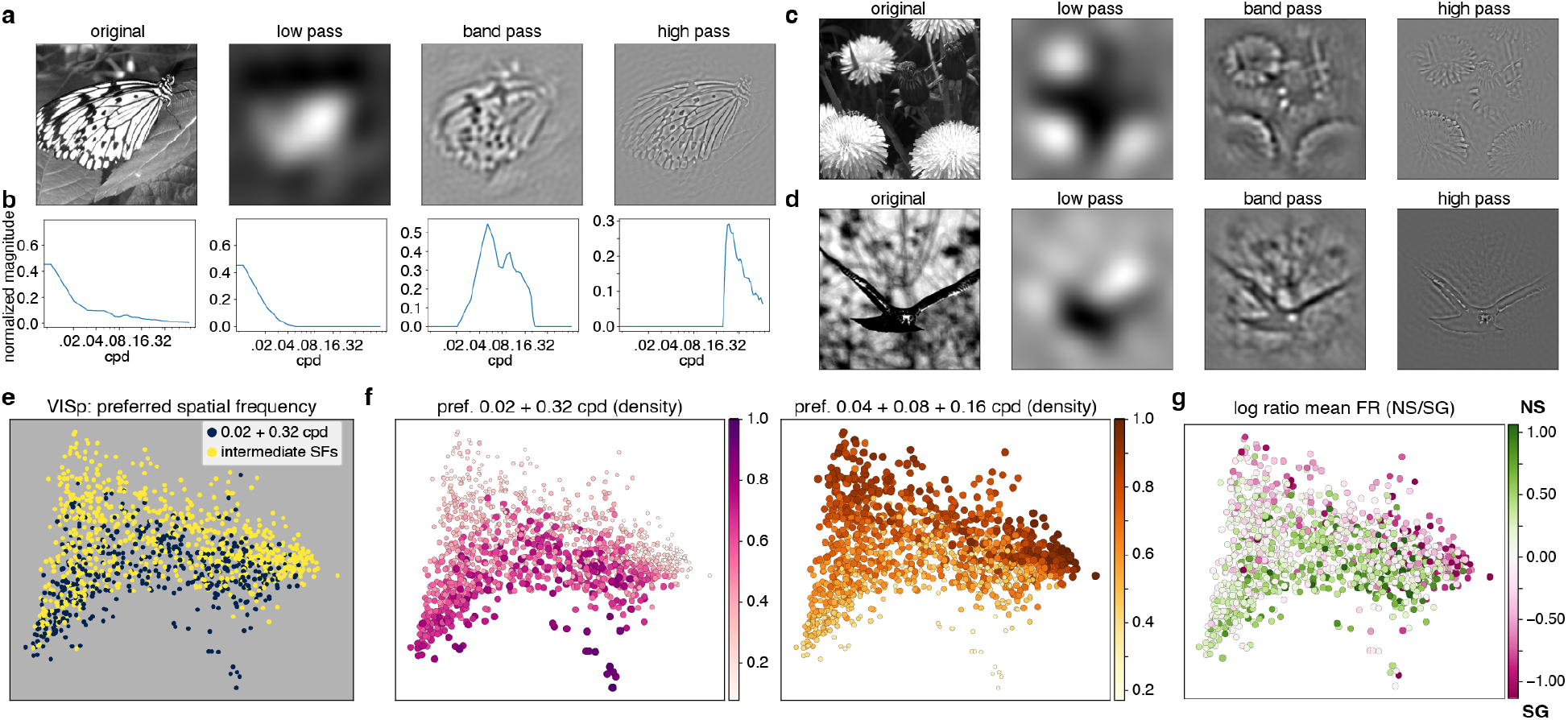
Natural scene statistics and neuronal preference on the encoding manifold. **a**, An example natural scene image and its low-pass, intermediate (band-pass), and high-pass filtered versions, and **b** associated spectra. **c–d**, Two more natural scenes and their filtered versions. **e**, Encoding manifold for VISp colored by spatial frequency preferences of neurons (as Fig. 6a): Neurons are colored according to their spatial frequency preference for static grating stimuli, grouping 0.02+0.32 (extreme frequencies, in yellow) vs. 0.04+0.08+0.16 (intermediate frequencies, in blue). **f**, Spatial frequency preferences summarized by the density over the data graph (see Methods) for extreme frequencies (left) and for intermediate frequencies (right). **g**, The log ratios from Fig. 4e, shown here for comparison with the distributions in **f**. The different spatial frequency bands thus provide insight into the U-shaped curves from Fig. 7b, suggesting a grouping of the low and high frequency content signals separated from the intermediate ones. For comparable plots in the higher visual areas, see Fig. S13.

Classically, natural scene analysis has been thought to proceed by a coarse-to-fine visual processing [30]. Object perception, figure/ground discrimination and hunting behavior [31, 62] are complex tasks, implicating computations across multiple visual areas [33], including feedback within and between them [27, 29] and recurrence [38, 74]. A recent study [61] found direct low– and high-frequency dynamics in natural scene viewing, extending classical results on dynamical frequency tuning [2, 13, 49, 69]. Since the encoding manifold considers the entire time course of responses, it is able to capture these dynamics—a possible signature of coarse-to-fine processing [46, 30]—in the manifold inference process. Perhaps while building biologically-motivated perceptual systems, following this study, one should consider coarse– *and* – fine processing.

### Encoding Manifolds

Encoding manifolds capture similarity in the responses over time of neurons exposed to a wide range of stimuli. They display neurons in relation to as many different stimulus features as one is able to illustrate. No other procedure of which we are aware lets one simultaneously visualize how the neurons that respond particularly well to one stimulus respond to many others. They disclose visually the relationships between, for example in the present data: firing rate, selectivity for stimulus orientation, spatial and temporal frequencies, cortical laminae, electrical waveform (related to excitatory versus inhibitory classes), overall selectivity (stimulus entropy), and preference for gratings versus natural scenes. This property should make encoding manifolds widely useful across biology when many relationships may be present but could go unnoticed.

## Methods

### Dataset

The “Neuropixels” single neuron recording dataset shared by the Allen Institute includes recordings from many cortical visual areas in alert wild type (C57BL6/J) mice [29]. Recording sites were localized to specific cortical areas by reference to intrinsic signal imaging maps and to cortical layer by transformation into the Allen Common Coordinate Framework. Details of all procedures are described at links from [4].

To summarize, visual stimuli were were displayed on an LCD monitor at a resolution of 1920 *×* 1200 pixels at 60 Hz refresh rate. Stimuli were presented monocularly, and the monitor spanned 120*^◦^ ×* 95*^◦^* of visual space prior to stimulus warping. Each monitor was gamma corrected and had a mean luminance of 50 cd/m^2^. To account for the close viewing angle of the mouse, a spherical warping was applied to all stimuli to ensure that the apparent size, speed, and spatial frequency were constant across the monitor as seen from the mouse’s perspective.

In our experiments, we analyzed responses to a subset of their “Brain Observatory 1.1” stimulus set, including 40 different drifting gratings, 120 stationary gratings, and 118 natural scenes [3]. We used data from all experiments that included these stimuli (32 mice in total). The drifting gratings were shown at a spatial frequency of 0.04 cycle/deg, 80% contrast, 8 directions of motion (0, 45, 90, 135, 180, 225, 270, and 315 degrees) and 5 temporal frequencies (1, 2, 4, 8, and 15 Hz), with 15 repeats per condition. The stationary grating set, also presented at 80% contrast, consisted of 120 gratings of 6 different orientations (0, 30, 60, 90, 120, and 150 degrees, clockwise from 0 = vertical), 5 spatial frequencies (0.02, 0.04, 0.08, 0.16, 0.32 cycle/degree), and 4 phases (0, 0.25, 0.5, 0.75), each presented for 0.25 seconds with no intervening gray period. The natural scene stimuli consisted of 118 natural images taken from the Berkeley Segmentation Dataset [48], the van Hateren Natural Image Dataset [67], and the McGill Calibrated Colour Image Database [57]. The images were presented in grayscale and were contrast normalized and resized to 1174 *×* 918 pixels. The images were presented in a random order for 0.25 seconds each, with no intervening gray period.

### Data preprocessing

As in Dyballa et al. [24], responses of a neuron to each stimulus (PSTHs) were kernel-smoothed using the Improved Sheather-Jones algorithm [12], using a bandwidth of 25 ms (typical value obtained for all visual areas when using automatic bandwidth selection) for drifting gratings and 10 ms for static gratings and natural scenes (between 39 and 50 trials). The `KDEpy` Python implementation [55] was used.

Responses to the drifting gratings were tested for statistical significance by performing a two-tailed Mann-Whitney test comparing the mean activity during the second half of the ISI immediately preceding the stimulus versus mean activity over any interval of the same length within the stimulus presentation. This strategy allowed transient responses to be able to reach the same level of significance as sustained responses. Responses were deemed significant if they had a *p*-value of at most 0.001. Only neurons that had a significant response to at least one variation of drifting grating were included in the data tensor (see below). The resulting number of neurons used were: *n* = 1261 for VISp, *n* = 910 for VISam, *n* = 961 for VISal, *n* = 525 for VISpm, and *n* = 810 for VISrl.

We were able to build meaningful encoding manifolds for gratings and flow stimuli in previous studies because these stimuli were globally homogeneous, so every neuron was simultaneously stimulated by similar visual features across time. This allows us to build meaningful tensors and generate a meaningful manifold embedding. Each natural scene, however, is really a different stimulus for different neurons depending on their receptive field positions; so aligning the same natural scene to all neurons in the tensor would not be feasible.

## Data analysis

### Neural encoding manifolds

Data tensors were constructed from the spikes detected in VISp and each of 5 higher cortical visual areas. Neural encoding manifolds were computed from those by using the procedure described in Dyballa et al. [24]. Briefly, it involves a dimensionality reduction pipeline consisting of two main steps: applying a permuted non-negative tensor factorization (pNTF) to a tensor built from the temporal responses of each neuron to each variation of grating stimuli, followed by manifold inference using iterated adaptive neighborhoods (IAN) [22] in combination with diffusion maps [18] to produce an embedding in which each point is a neuron. We briefly overview this material here for completeness; please refer to Dyballa et al. [24] for details and further discussion. Code is available at https://github.com/ `dyballa/NeuralEncodingManifolds`.

### Tensor decomposition

We organize the data as a *3-way tensor*, NEURONS^(1)^*×* STIMULI^(2)^*×* RESPONSE IN TIME^(3)^. (For an introduction to tensors, see [72].) In standard notation [40]:

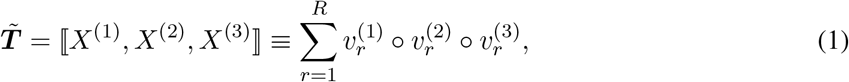

where *◦* denotes vector outer product and the *factors* are collected into the factor matrices *X*^(*k*)^, with individual factors 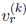 as columns. A *component* is an associated set of factors, one from each tensor mode.

Factors are normalized by computing 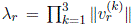, where *r* indexes the component and *k* the factor mode. Collecting these scalars into a vector *λ ∈* R*^R^*, we have:

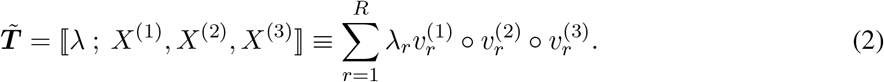

Since neuronal firing rates are non-negative, we adopt a non-negative tensor factorization (NTF) algorithm [14] based on CP decomposition. This minimizes the squared reconstruction error [16]:

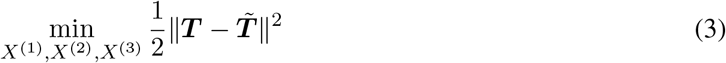

subject to the non-negativity constraint *X*^(^*^k^*^)^ *≥* 0*, k ∈* [1, 2, 3] [42, 16]. Non-negative tensor factorization is further advantageous in the sense that it imposes a part-like decomposition [42, 16].

For implementation, we use a gradient-based direct optimization approach (OPT) [1] in the Tensor Toolbox [7], modified to allow permutation of the response factors. This allowed us to exploit data limitations, since two narrowly-tuned simple cells (for example) are similar except in their preferred orientations, effectively collapsing the orientation-preference coordinate on the manifolds.

Because drifting and static gratings have different trial lengths and a different number of stimulus variations, two tensors were constructed for each cortical area: one containing the drifting grating responses (in which the stimulus mode consisted of the 5 temporal frequencies) and another for the static grating responses (in which the stimulus mode consisted of the 5 spatial frequencies). The third (response) mode consisted of the vectorized peri-stimulus time histograms obtained by concatenating the responses to all directions (8, for drifting gratings) or orientations (6, for static gratings at the preferred spatial phase). Thus, the drifting-grating tensor had dimensions neurons *×* temporal frequencies *×* (directions *×* time), while the static-grating tensor had dimensions neurons *×* spatial frequencies *×* (orientations *×* time).

Importantly, orientation (for static gratings) and direction (for drifting gratings) were incorporated into the temporal mode rather than treated as independent stimulus modes. This design allows the permuted tensor decomposition to normalize away the particular preferred orientation or direction of individual neurons, so that neurons with similar tuning structure but different preferred orientations can be treated as functionally similar. By contrast, spatial and temporal frequencies were retained as explicit stimulus variables because their organization across the neuronal population was a primary object of study. Consequently, the manifold is designed to be insensitive to orientation/direction preference itself, while remaining sensitive to properties such as tuning sharpness, response dynamics, and frequency selectivity.

Finally, we also built a tensor combining all visual areas. An equal number of neurons (*n* = 500) were sampled uniformly at random from each area, for a total of *n* = 3000. All additional steps were kept the same as those for building the manifolds for individual areas. The results are shown in Fig. S12.

### Neural encoding space

The first mode of the tensor decomposition consists of the neural factors, which together form the neural matrix *N* (factors as columns). Letting Λ denote the diagonal matrix containing the component magnitudes *λ_r_*, the reconstruction of the tensor can be written as

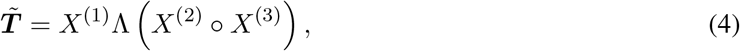

where the outer product between the stimulus and response factors of each component forms a vectorized stimulus-response basis. The matricized tensor with respect to the neural mode can therefore be expressed as

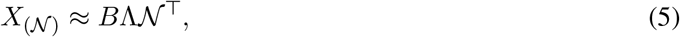

where each column of *B* is obtained by vectorizing the outer product of the corresponding stimulus and response factors. Thus, each neuron is represented as a linear combination of these stimulus-response vectors. We define the scaled neural matrix

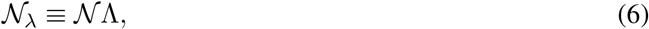

whose rows provide a low-dimensional representation of the neurons in the neural encoding space.

The number of tensor components was selected using the normalized explained-variance criterion introduced in [24]. Briefly, candidate decompositions with different numbers of components were computed, and principal component analysis was applied to the corresponding neural matrices. The number of tensor components was chosen as the smallest value for which the normalized explained variance of the neural matrix reached a plateau, thereby avoiding underfactoring while also avoiding overfactoring, which tends to split meaningful components or model noise. As in [24], multiple random initializations were evaluated for each candidate number of components, and the solution with the lowest reconstruction error was retained.

For each visual area, separate neural matrices were obtained from the drifting-grating and static-grating tensor decompositions. These matrices were concatenated column-wise to form a single neural representation for each neuron. Principal component analysis (PCA) was then applied to the combined matrix. The minimum number of principal components required to explain at least 80% of the variance was retained for manifold inference.

### Similarity kernel and diffusion maps

The data graph for manifold inference is built from pairwise distances computed from the orthogonal PCA representation described above. Since neighborhoods may vary, we use the IAN kernel [22], a multiscale Gaussian kernel in which the neighborhood scales are automatically inferred from the local geometry of the data in a (relaxed) optimal fashion. The IAN weighted graph yields the similarity matrix on which the diffusion maps algorithm [18, 17] is based. Diffusion maps are a non-linear spectral embedding method based on the eigendecomposition of the normalized graph Laplacian that tends to better preserve the topology of the data [22]. The resulting diffusion coordinates are then used to embed the neurons in (reduced) stimulus-response coordinates (see above). The standard parameters *α* = 1 (Laplace-Beltrami approximation) and diffusion time *t* = 1 were used throughout. For some manifolds, a minority of points that were too far from their nearest neighbor in the embedding were treated as outliers and removed from the analysis (9 from VISam, 3 from VISpm, 2 from VISlm, and 1 from VISrl).

One of the main advantages of using a spectral embedding method such as diffusion maps is the natural way in which multi-dimensional aspects of the data can be visualized by looking through the various ‘diffusion coordinates’ produced (denoted as ‘dc’s in the figures). Importantly, different diffusion coordinates concentrate different aspects of the data. Other popular methods, such as t-SNE [66] and UMAP [50], approach this differently. They are based on a gradient descent optimization that focuses on preserving nearest neighbors at the expense of global structure. Both t-SNE and UMAP tend to collapse these multiple dimensions in ways that end up stochastically mixing together the global coordinates that run through the manifold geometry (see Fig. 29 in [22] for an example of this phenomenon). Diffusion maps, by contrast, yield deterministic results.

Our figures show a projection of the diffusion maps into two dimensions, while the intrinsic dimensionality (the NCD algorithm was used, see [22] for computation details) was between 5 and 6 for all cortical areas; see Fig. S3. Just as in principal component analysis, where different coordinates (principal components) emphasize different aspects of the data, so too do lower-dimensional projections of diffusion maps illustrate different features of organization. We chose the two-dimensional projections of the manifolds shown in the figures to emphasize how they organize the properties, such as orientation selectivity, noted in the text. In particular, we selected the pair of diffusion coordinates that maximized the *coefficient of determination* (*R*^2^) for a plane fit to the function defined by the label values (e.g., OSI or electrical waveform type density) over the domain given by the 2-dimensional coordinate values for all data points. To illustrate this for VISp, we fit a plane in diffusion coordinates dc0 and dc3 to OSI values resulting in *R*^2^ = 0.99; to narrow-spiking cell densities resulting in *R*^2^ = 0.68 and to broad-spiking cell densities resulting in *R*^2^ = 0.70. This provides evidence for the success of the method in organizing cell properties in an unsupervised manner, since those labels were not used to compute the manifolds. Other coordinates organize different properties, such as temporal frequency preference (Fig. S2).

### Density of categorical properties over the manifold

The local density of categorical properties around each neuron in a manifold (such as layer, electrical waveform type, and preferred spatial frequency) was computed as the fraction of adjacent nodes in the non-weighted IAN graph. Some densities as displayed as non-negative z-scores (computed from the global distribution) for a more clear visualization, highlighting the regions where in which the property was most present.

### Natural scene vs. static grating selectivity ratio

Selectivity ratios for individual neurons were computed as the natural logarithm of the ratio between their firing rate to natural scenes and their firing rate to static gratings. Thus a selectivity ratio of 0 meant the neuron produced exactly the same magnitude of response to both stimulus classes; and a positive (respectively, negative) value indicated a stronger response to natural scenes (resp., static gratings). The magnitudes of responses to these two stimulus classes are readily comparable because both stimuli consist of static images presented during the same time interval of 250 ms.

Following the methodology from the Allen Brain Observatory SDK [5], mean firing rates (FR) for each stimulus class (namely, drifting gratings, static gratings, and natural scenes) were computed as the mean number of spikes per second of stimulus presentation across all trials of the same stimulus, kernel-averaged across all stimulus variations within the same class. For static gratings, ‘variations’ meant different orientation (6), spatial frequency (5), and phase (4); for natural scenes, ‘variations’ meant each of 118 scenes used. We also computed the maximum FR for a given stimulus class as the maximum kernel-averaged FR across all stimulus variations (in other words, the FR to the preferred stimulus variation). Thus, two ratios were computed: one using the ‘mean’ responses (“mean FR ratios”), and another using the ‘maximum’ responses (“max FR ratios”). As a complementary analysis, we also computed population mean FR to natural scenes and static gratings separately as a function of preferred spatial frequency (Fig. S18).

To assess whether the observed U-shaped relationship between preferred spatial frequency and the ratio of responses to natural scenes versus static gratings could arise by chance, we performed a permutation test. For each visual area, natural-scene firing rates were randomly shuffled across neurons while static-grating responses and preferred spatial-frequency labels were left unchanged. For each permutation, we recomputed the spatial-frequency-dependent ratio curve and quantified its U-shape using the statistic

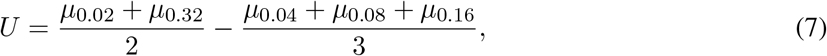

where *µ* denotes the mean log(NS/SG) ratio of neurons preferring the indicated spatial frequency. Positive values indicate larger natural-scene preference among neurons preferring extreme spatial frequencies relative to intermediate spatial frequencies. A null distribution was obtained from 10,000 independent permutations, and permutation-test p-values were computed as the fraction of null U-statistics greater than or equal to the observed value (see Fig. S14).

For the layer-specific analyses shown in Fig. 8, one-sample t-tests against zero were performed on the distributions of log response ratios.

### Additional metrics

We used precomputed stimulus metrics available from the Allen Brain Observatory SDK [5] for obtaining firing rate, global orientation selectivity index (OSI), preferred spatial frequency (drifting gratings), and preferred phase (static gratings) for each unit. Receptive field position and area estimation followed the Allen SDK methodology of using response maps to localized Gabor stimuli and fitting those to a 2-D Gaussian distribution. Laminar position and mean electrical waveform were also obtained directly from the Allen SDK. Individual neurons were classified into broad spiking (putative excitatory) (VISp *n*=944, VISpm *n*=397, VISal *n*=726, VISam *n*=669, VISrl *n*=634, VISlm *n*=487) or narrow spiking (putative inhibitory) (VISp *n*=306, VISpm *n*=124, VISal *n*=227, VISam *n*=236, VISrl *n*=171, VISlm *n*=110) based on their extracellular spike waveform (see [54]). In addition, a stimulus entropy index (Fig. 2c) was defined as 2*^H^*, where *H* is the base-2 entropy of the vector containing the relative response magnitudes (divided by their sum) of a neuron to the 5 temporal frequencies used in the drifting gratings experiments. It therefore ranges between 0 (case in which the neuron responds to a single stimulus) and 5 (when it responds with uniform magnitude to all stimuli).

### Spatial-frequency analysis of stimulus ensembles

To compare the spatial-frequency content of the natural-scene and static-grating stimulus ensembles, each image was first mean-centered and multiplied by a separable two-dimensional Hann window to reduce edge discontinuities and spectral leakage. A two-dimensional Fourier magnitude spectrum was then computed using the FFT module in the `Numpy` Python library [28]. The azimuthally averaged radial profile of the two-dimensional Fourier magnitude spectrum was computed for each image and then averaged across all stimulus instances within a given stimulus class (118 natural scenes and 120 static gratings). Similarity between the resulting average radial spectra was quantified using Pearson correlation and cosine similarity.

### Natural scene filtering

Filtered versions of the natural scenes were created by first computing fast Fourier transforms of each image as described above. For the low-pass filtered version, the Fourier domain was cropped with a disk of radius 0.04 cpd; for the band-pass version, an annulus with inner radius 0.04 cpd and outer radius 0.22 cpd; and for the high-pass version a disk with radius 0.22 cpd (see example in Fig. S17).

## Acknowledgements

Research supported by NIH Grant EY031059, NSF CRCNS Grant 1822598, the European Commission’s Marie Skłodowska-Curie Action Grant agreement no. 101207931 (LD), and an unrestricted fund to the Jules Stein Eye Institute from Research to Prevent Blindness and NIH P30 – EY000331 (GDF). We thank the Allen Institute for the use of their data and Cris Neill for discussions.

---

**Fig. S1:**
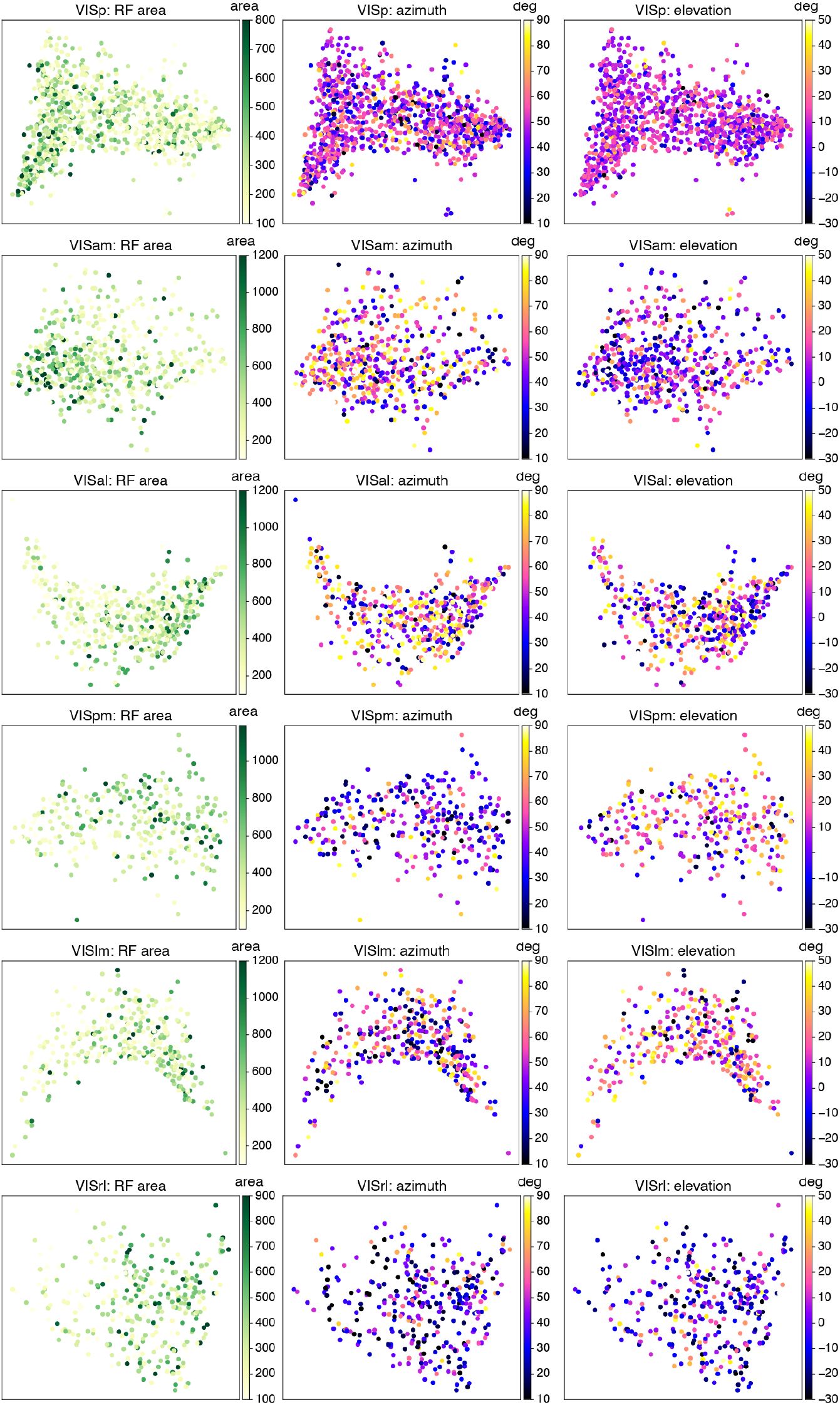
Receptive field properties show no clear concentrations on the manifolds. Manifolds colored by each neuron’s receptive field (RF) area (left), and receptive field position in the visual field: azimuth (middle) and elevation (right), in degrees of visual angle. A lack of clear concentrations was observed for all visual areas.

**Fig. S2:**
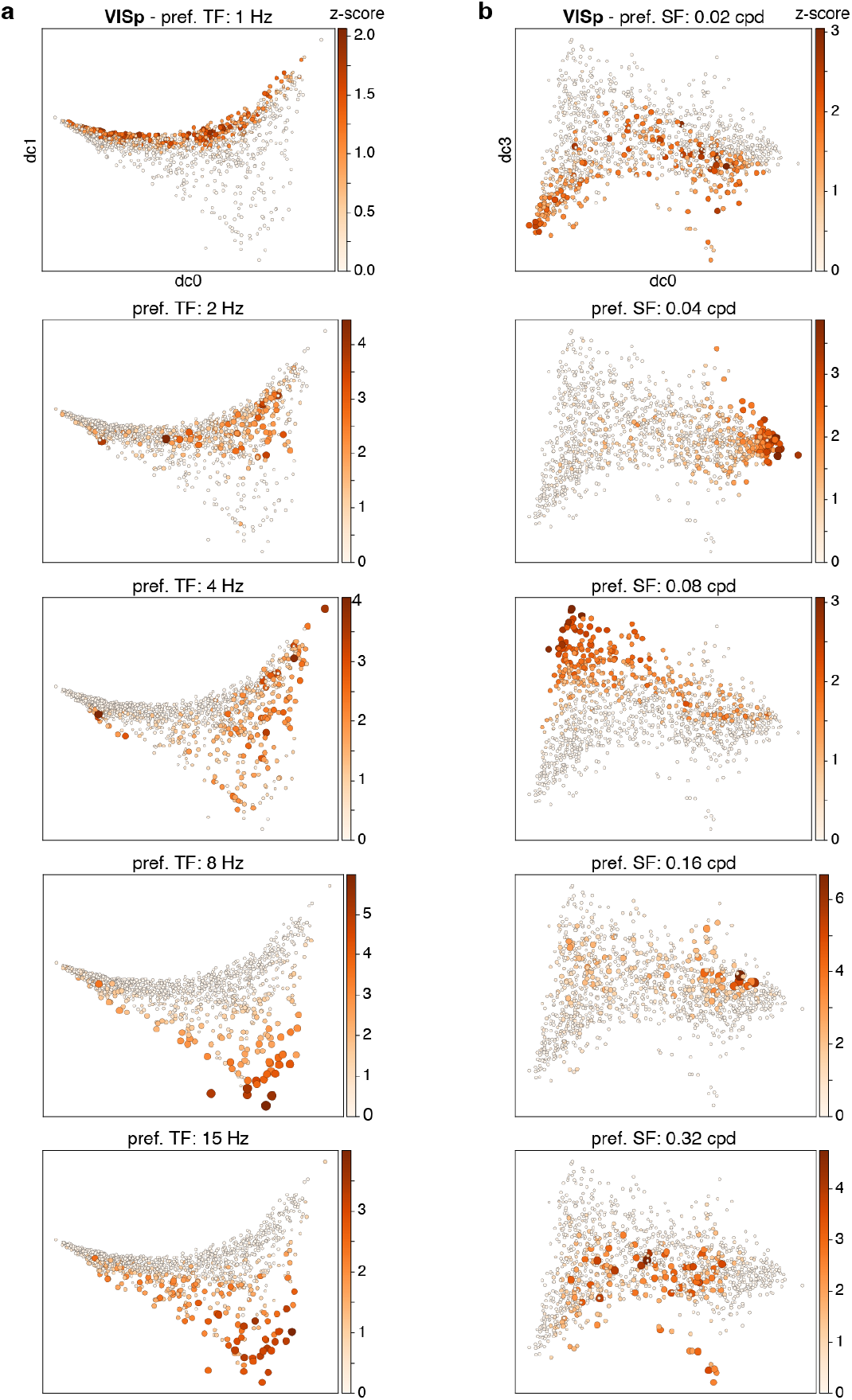
Temporal and spatial preferences across the VISp manifold. Each manifold is colored by the local density (z-scores, see Methods) of cells preferring a particular temporal (**a**) or spatial (**b**) frequency from the grating stimuli. **a**, Temporal frequency preference is organized by a gradient along the diffusion coordinate ‘dc1’: preference for lower frequencies concentrate near the top, higher frequencies at the bottom. **b**, In contrast, spatial frequencies are not organized sequentially: notice how along diffusion coordinate dc3 preference for 0.02 cpd and 0.32 cpd concentrate mostly towards the bottom, while the intermediate preferences (0.04, 0.08, and 0.16) concentrate in the upper portion. This trend is summarized in Fig. 6 and compared to the distribution of preference for gratings vs. natural scenes. The same is shown for the higher visual areas in Fig. S13.

**Fig. S3:**
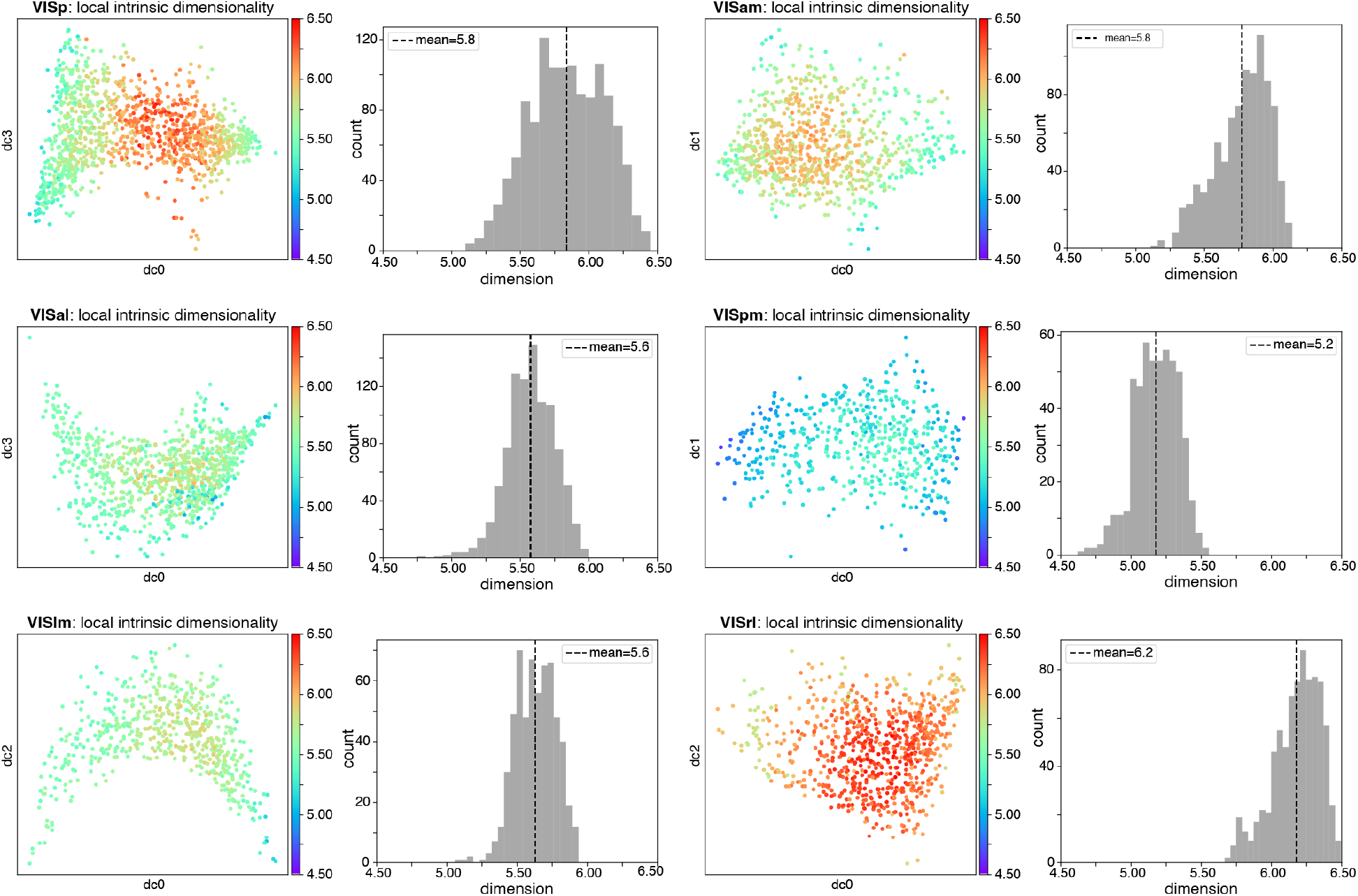
Local intrinsic dimensionality. All areas had a roughly homogeneous intrinsic dimensionality (ID) throughout their manifolds, with no particular feature in the distribution of dimensionality that stood out (e.g., no clear correlation with any of the properties studied). The mean ID varied between 5.6 and 5.8 for all areas except for VISpm (slightly lower, mean = 5.2) and for VISrl (slightly higher, mean = 6.2). VISp also had a large portion of it going slightly beyond 6.

**Fig. S4:**
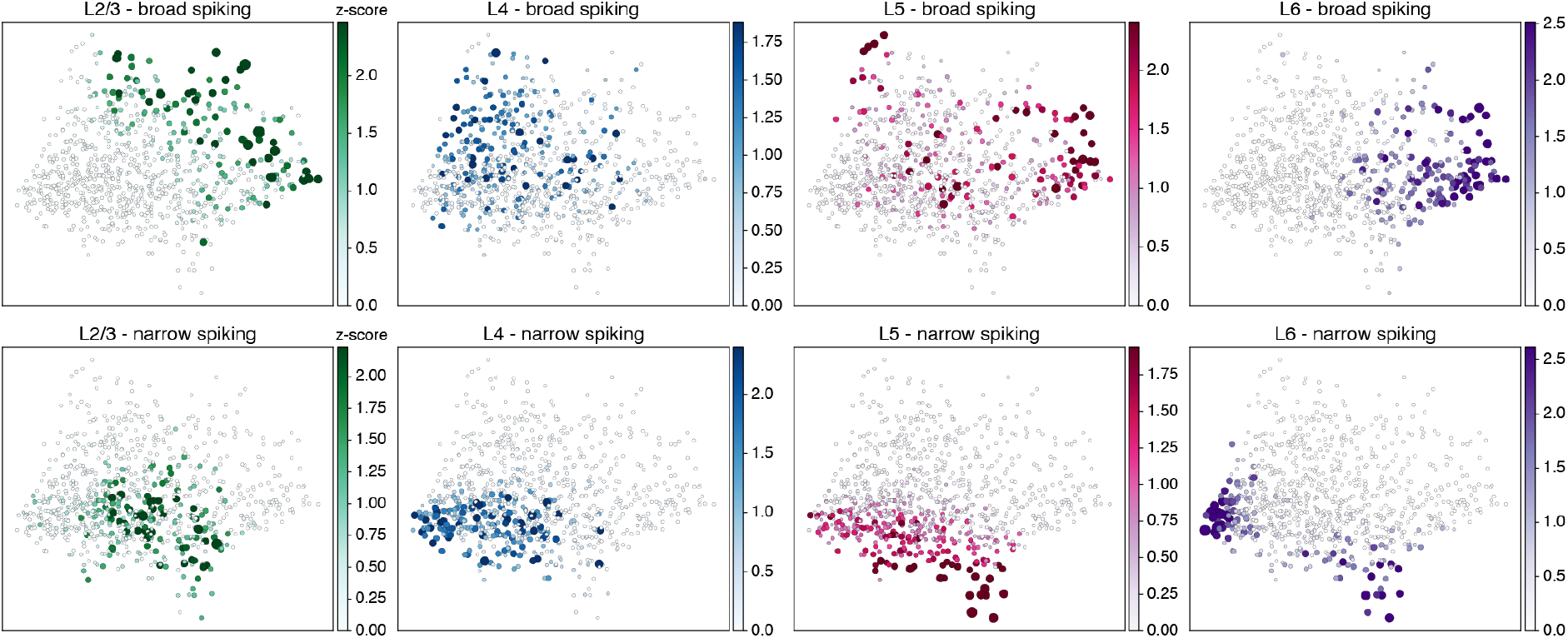
Laminar densities across the VISam manifold. Density of broad– and narrow-spiking neurons by layer. Note the distinctness of layer 2 vs. layers 5 and 6 for broad-spiking neurons, and the multi-lobed density of narrow-spiking neurons in layer 6.

**Fig. S5:**
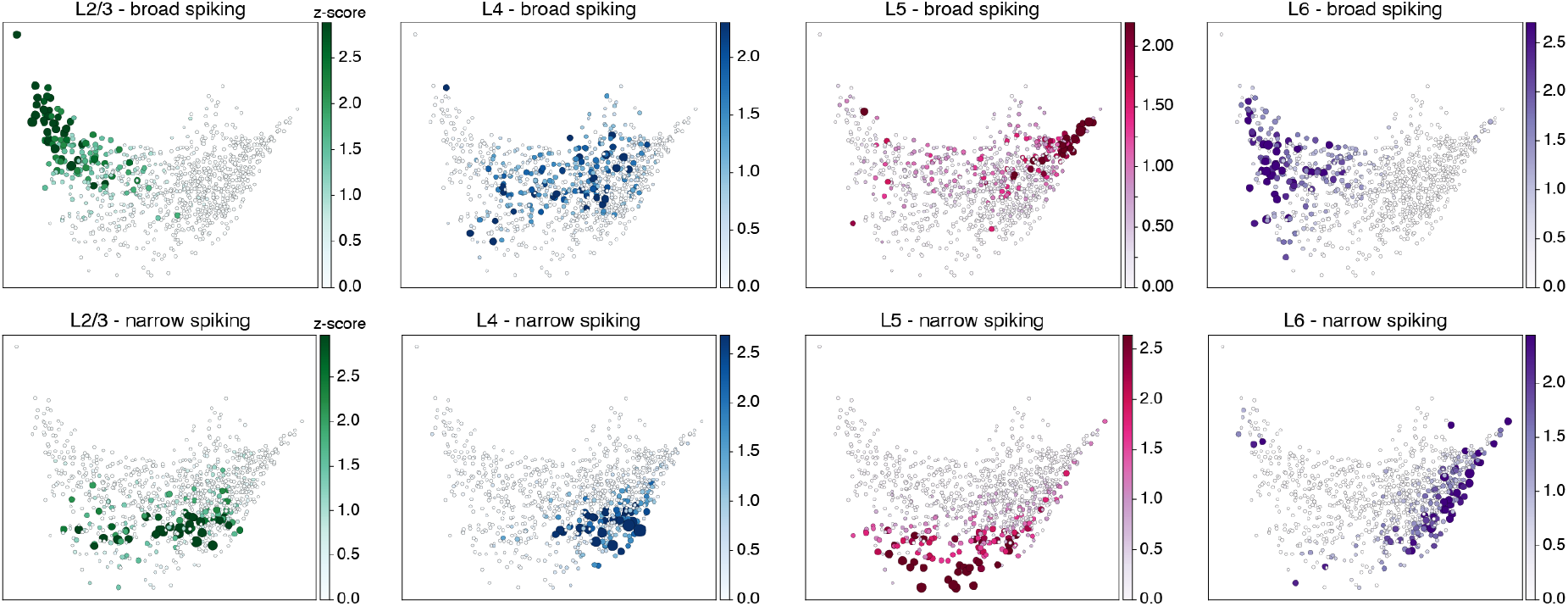
Laminar densities across the VISal manifold. The distributions of broad– and narrow-spiking neurons in the different cortical layers of VISal are strikingly diverse. In general, within any one layer, the concentrations of broad– and narrow-spiking neurons are largely but not fully complementary. Broad-spiking neurons in layer 6 are distributed over a region consisting mostly of neurons that prefer gratings over natural scenes, whereas the layer 6 narrow-spiking neurons are concentrated in the right-most portion of the manifold. Layer 2/3 is similar to layer 6. Layer 4’s broad-spiking neurons extend into the center of the manifold, differently from any of the other layers, and its distribution of narrow-spiking neurons largely intersects that of layer 2/3. Layer 5 in VISal has the most unusual distribution: broad-spiking neurons are most concentrated in the upper right, where we find a similar overall concentration of both broad– and narrow-spiking neurons (compare with Fig. S10). Layer 5’s narrow-spiking neurons occupy the lowest portion of the manifold, different than any other layer, indicating that they exhibit unique response properties among neurons in VISal. These relations between laminar distributions on the manifold are quite distinct from those found, for example, for VISpm (see Fig. S6, which can provide insights to the key functional differences among higher visual areas.

**Fig. S6:**
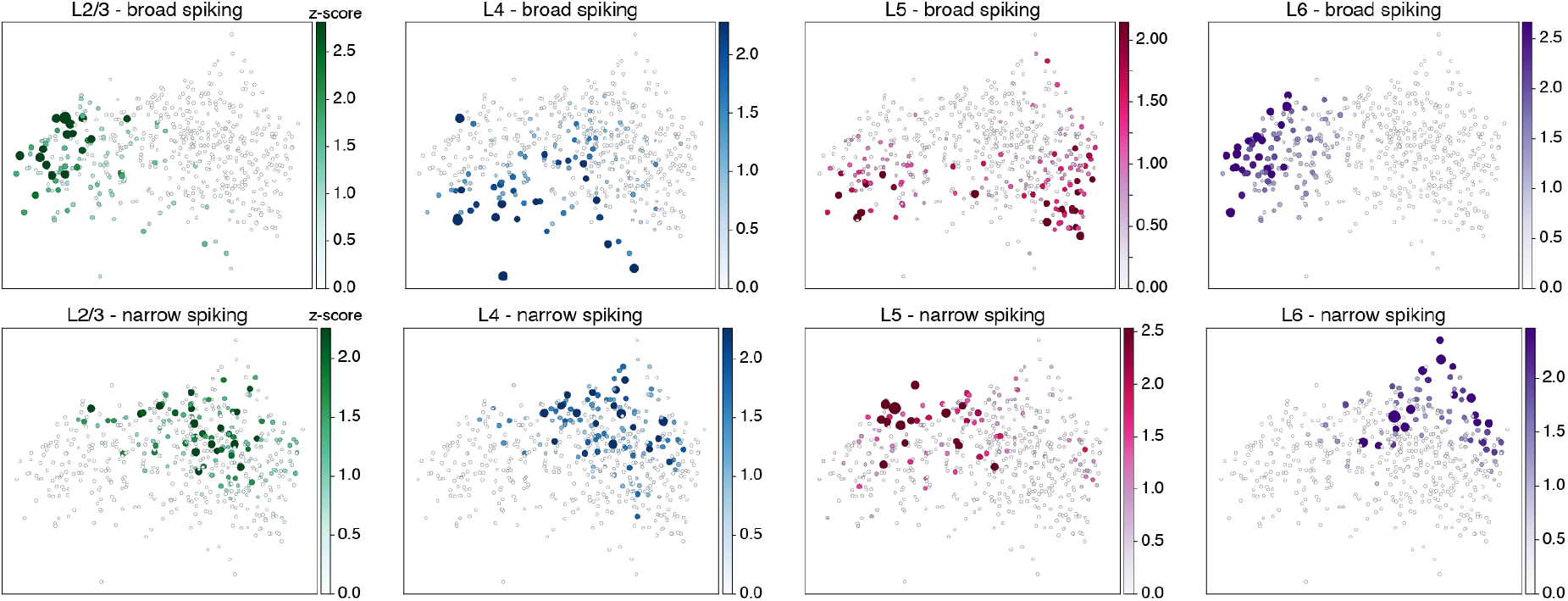
Laminar densities across the VISpm manifold. Layer 2/3 had significantly different concentrations of excitatory neurons from layer 4’s, and layer 5 showed two populations of excitatory neurons. Layer 6, on the other hand, was closer to layer 2/3 in distribution.

**Fig. S7:**
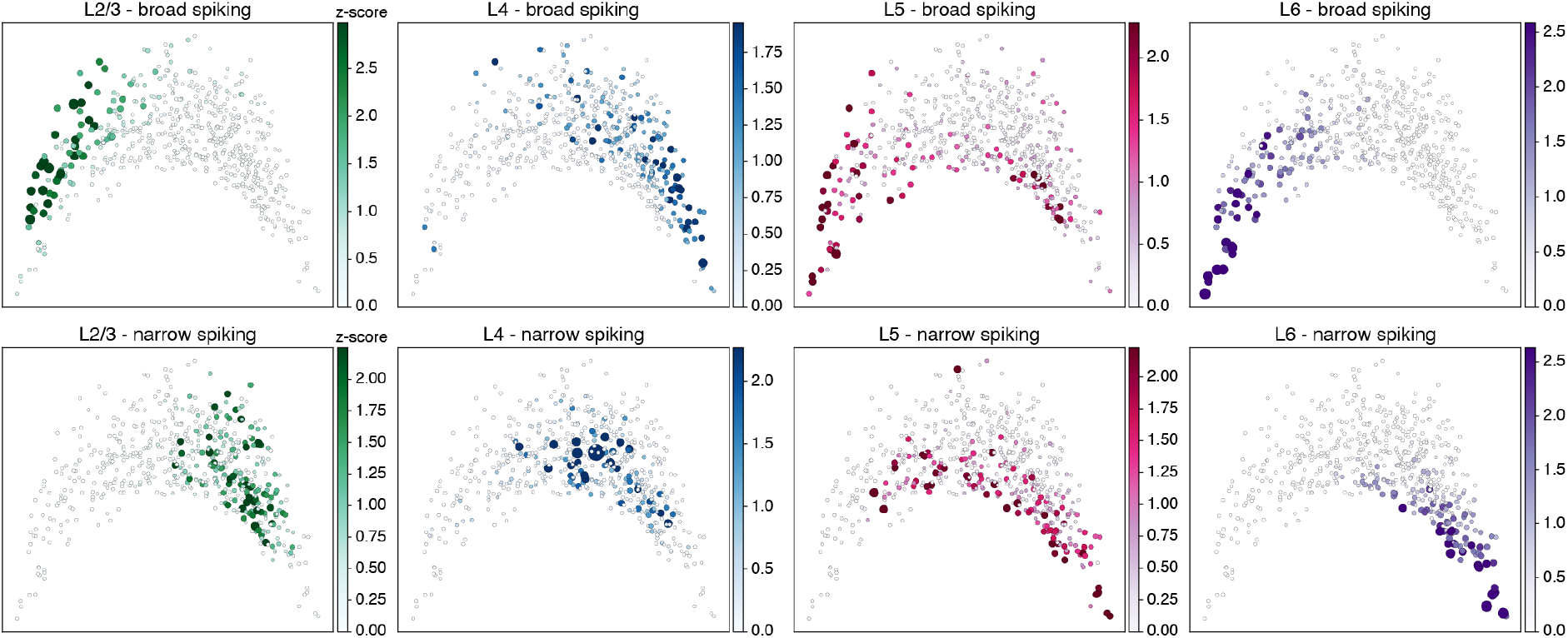
Laminar densities across the VISlm manifold. Density of broad– and narrow-spiking neurons by layer. Note the similarity of layers 2 and 6 for broad-spiking neurons, and the distinctness of layer 4 for narrow-spiking neurons.

**Fig. S8:**
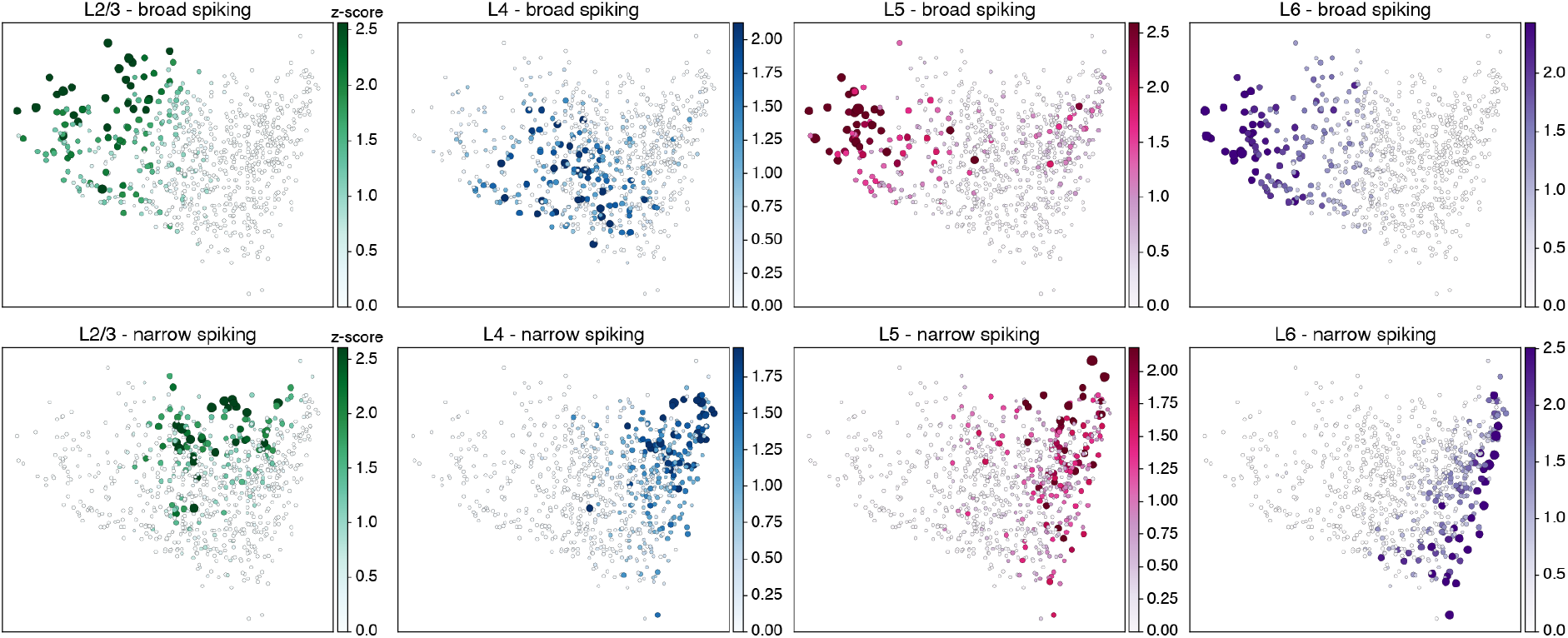
Laminar densities across the VISrl manifold. Like VISlm, there is a multi-lobed distribution of broad-spiking neurons in layer 5, and similar agreement between layers 2 and 6.

**Fig. S9:**
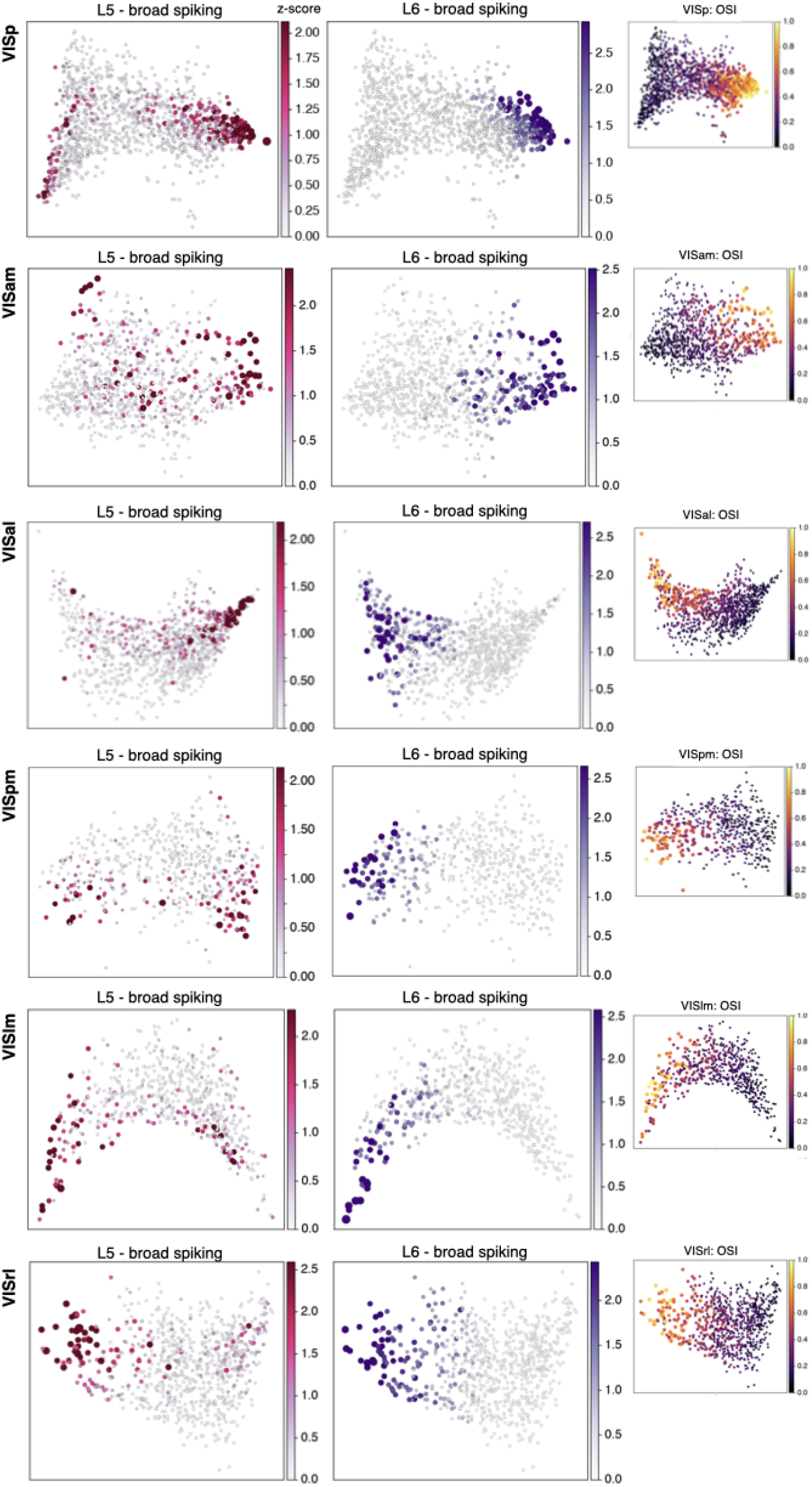
Relationship between distributions of layers 5 and 6 for broad-spiking neurons. In VISp, layer 5 (left column) exhibited two distinct concentrations, one of which overlapped almost completely with layer 6 (middle column) in the region of highest orientation selectivity (OSI, shown in inset). Layer 5 neurons in the VISal, VISpm, VISlm, and VISrl manifolds were also organized in two clusters, with high overlap of one cluster with the highly orientation-selective layer 6 neurons. On the other hand, in VISal layer 5 showed very small overlap with layer 6, even though the latter remained highly selective.

**Fig. S10:**
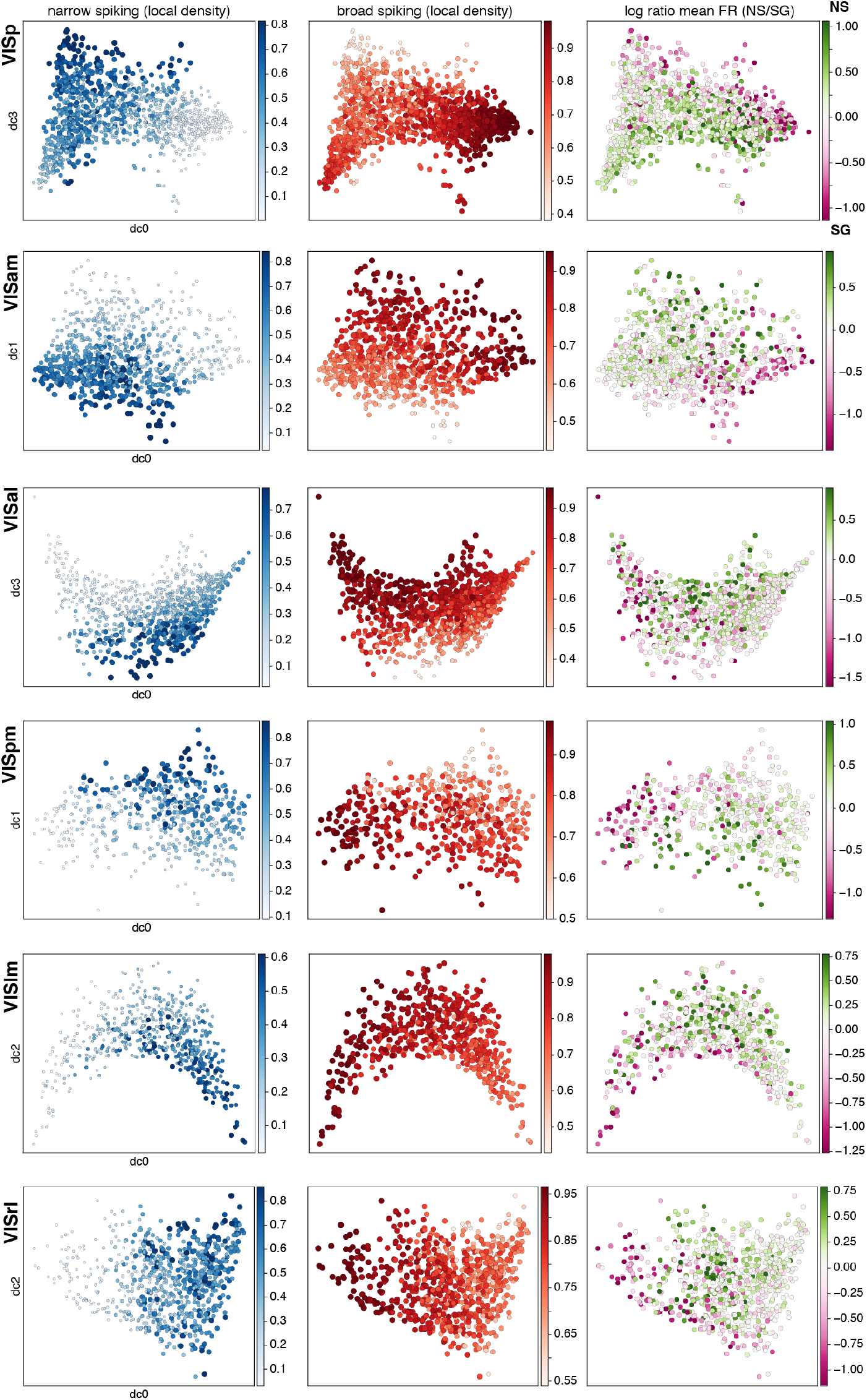
Smooth property distributions across all visual areas. **Left**, Encoding manifolds colored by the density of narrow-spiking and **middle**, broad-spiking neurons. **Right**, Encoding manifolds colored by the ratio of mean responses to natural scenes (NS) over static gratings (SG). In all areas, the manifolds shows nearly complementary densities for the two electric waveform shapes, even though firing rate information was not directly provided to the algorithm. Comparing the three columns reveals no clear overlap (implying no strong correlation) of a particular waveform shape with a preference for either gratings or natural scenes.

**Fig. S11:**
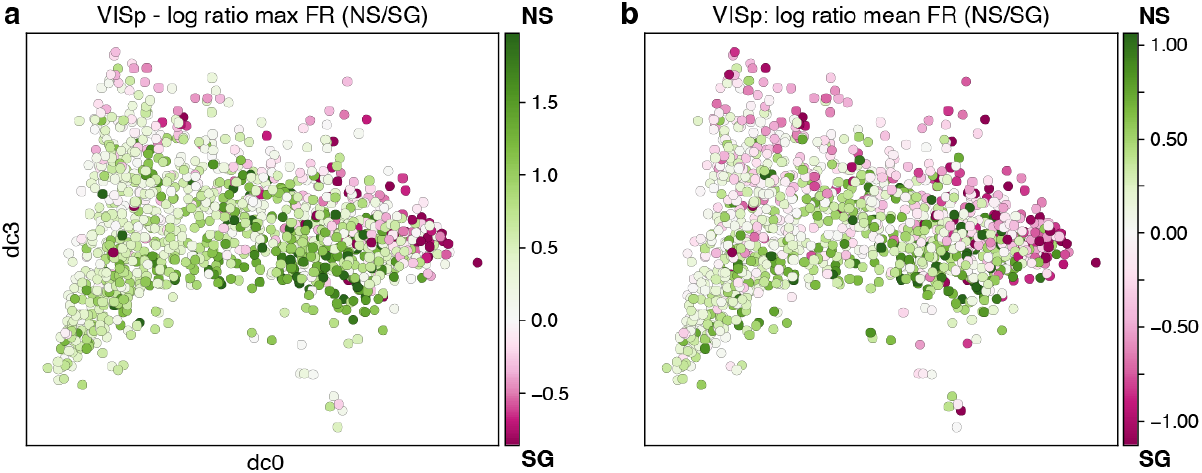
Ratio distribution is robust to maximum firing rate. **a**, Log ratios of the maximum firing rate over all natural scenes divided by the maximum firing rate over all static gratings are plotted on the VISp encoding manifold. **b**, For comparison, an analogous plot using mean firing rates when computing log ratios (cf. Fig. 4 in the main text).

**Fig. S12:**
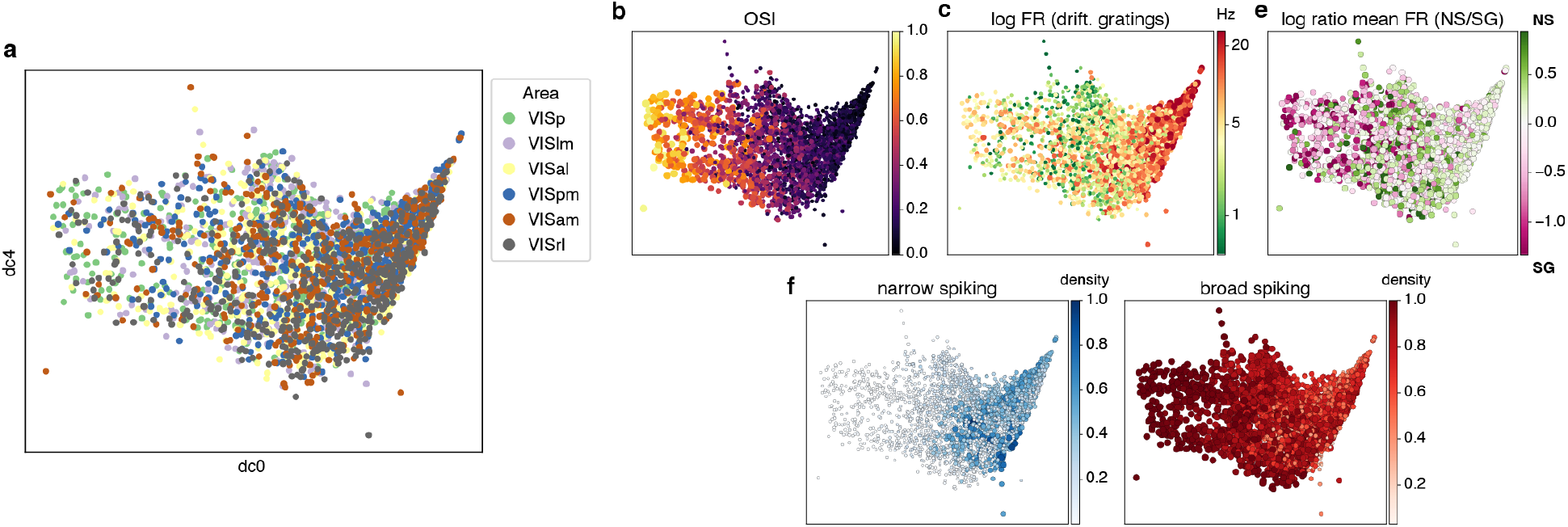
A manifold for all visual areas combined. After co-embedding all areas into a single encoding manifold, the 6 areas end up largely mixed. This is to be expected, since every area was organized for the gratings-based stimulus set around qualitatively similar coordinates (cf. Fig. 3 in the main text).

**Fig. S13:**
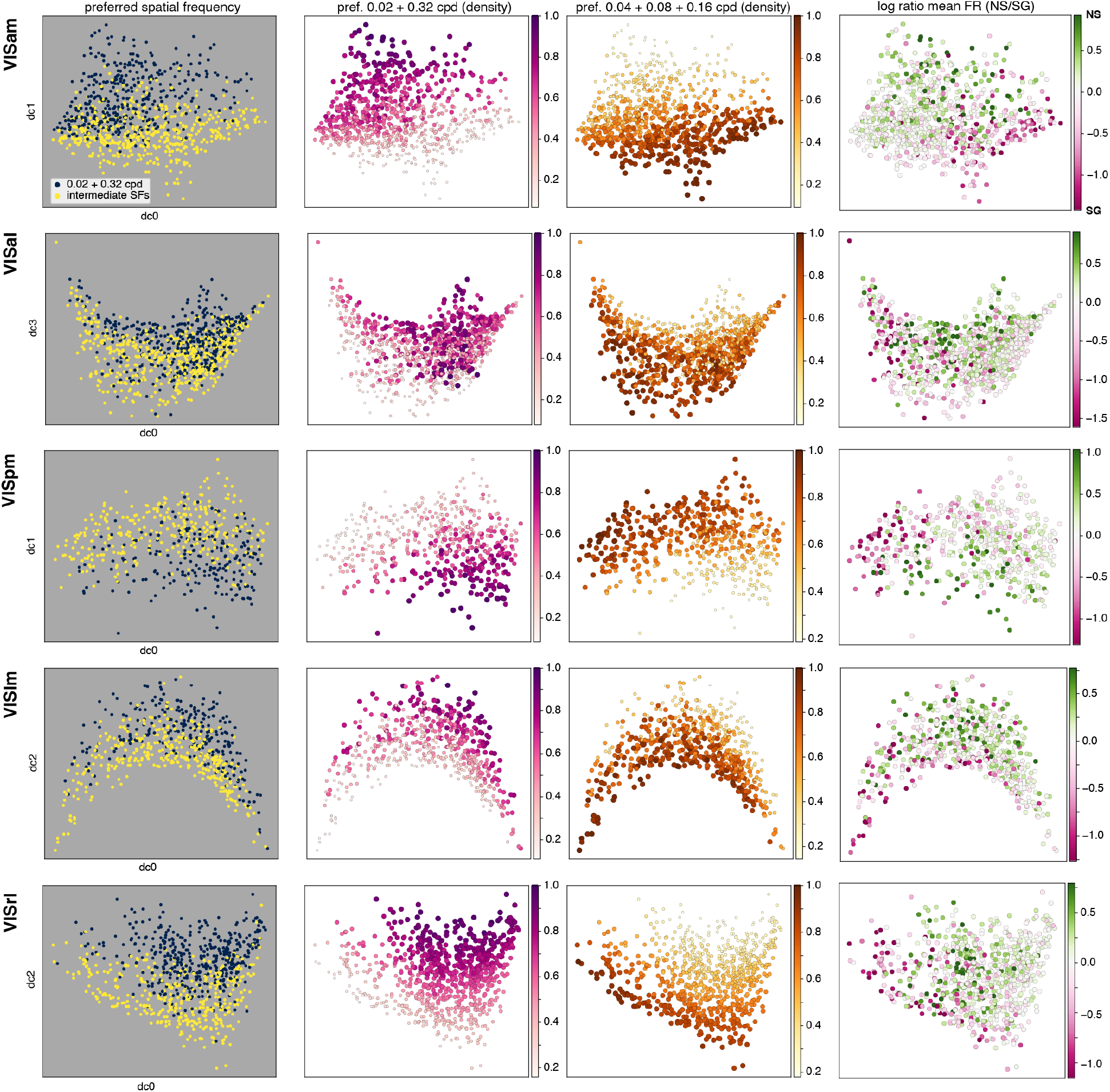
Preference for extreme vs. intermediate frequencies. A grid of colored manifolds, in which each row is a visual area and each column is (from left to right): preference for high and low spatial frequencies vs. intermediate ones; local density of cells preferring high and low spatial frequencies; local density of cells preferring intermediate spatial frequencies; and the log ratio of mean firing rate to natural scenes (NS) to static gratings (SG). Corresponding plots for VISp are shown in Fig. 6a–c in the main text.

**Fig. S14:**
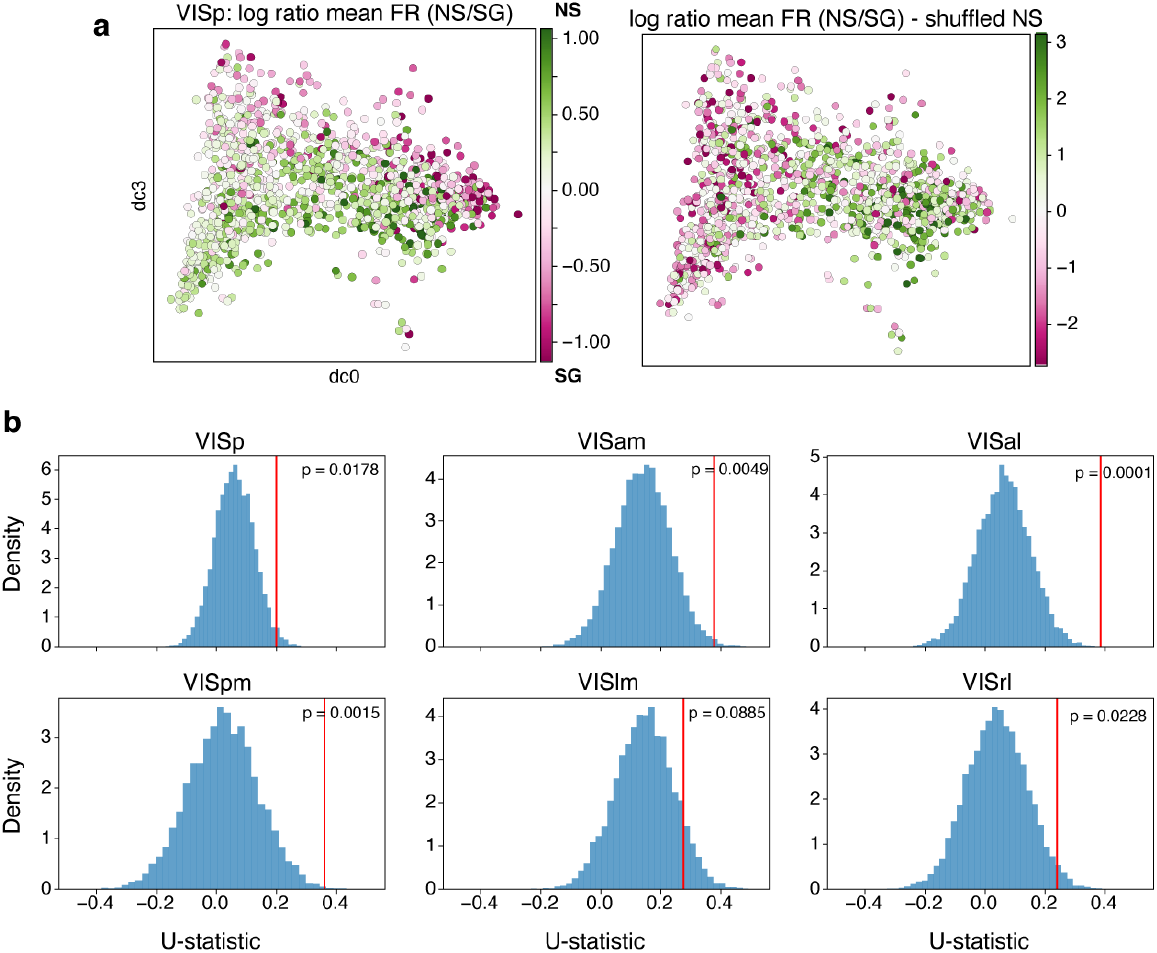
Statistical validation of relationship between preferred spatial frequency and natural-scene preference. To rule out the possibility that the distributions based on log ratios between natural scenes (NS) and static gratings (SG) are merely a consequence of the gradient in firing rates to static gratings, natural-scene firing rates were randomly shuffled across neurons while static-grating responses and preferred spatial-frequency labels were left unchanged. **a**, Comparison between the original (left) and shuffled (right) VISp manifold colored by the log ratio of mean responses to NS versus SG. Shuffling destroys the original organization, yielding an almost orthogonal trend since the resulting ratios are now driven primarily by SGs responses. The same was true of all other areas. **b**, Null distributions of the U-statistic (see Methods) obtained from 10,000 independent permutations for each visual area. Red lines indicate the observed U-statistics, and corresponding permutation-test p-values are shown in each panel. In five of the six visual areas studied, the observed U-statistic exceeded that expected under the permutation null distribution, supporting the conclusion that the observed U-shaped relationship is unlikely to arise from chance associations between preferred spatial frequency and natural-scene responses.

**Fig. S15:**
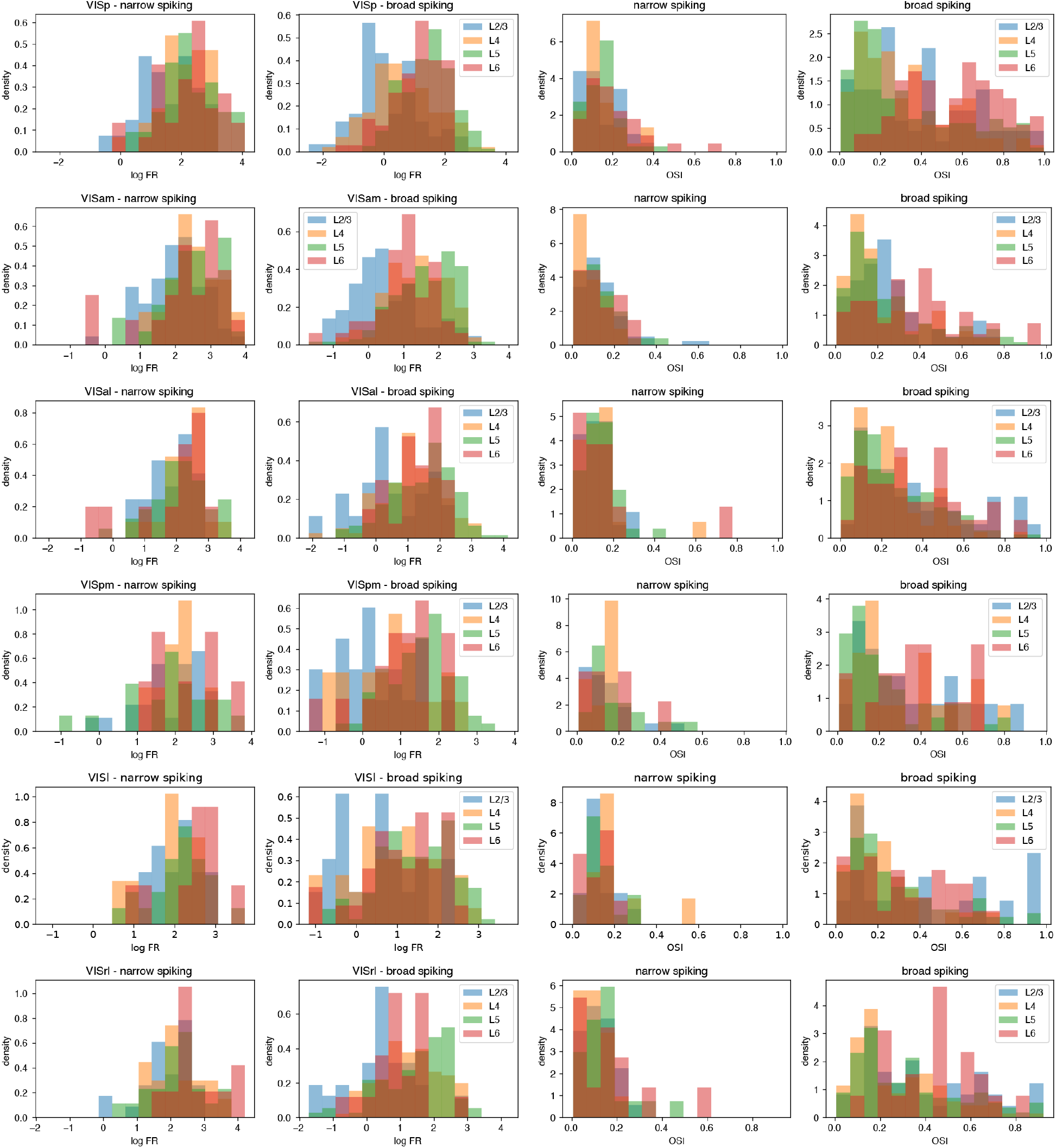
Laminar distribution analysis based on orientation selectivity and firing rates alone. A naive, histogram-based analysis of laminar distribution based on traditional cell properties such as firing rate (FR) and orientation selectivity index (OSI) does not reveal clear differences or similarities between visual areas (compare with Figure 2h,i and Figures S4, S5, S6, S7, and S8. This highlights the advantages of using data-driven, unsupervised approaches such as the encoding manifold method for understanding how functional properties organize across a large neuronal population.

**Fig. S16:**
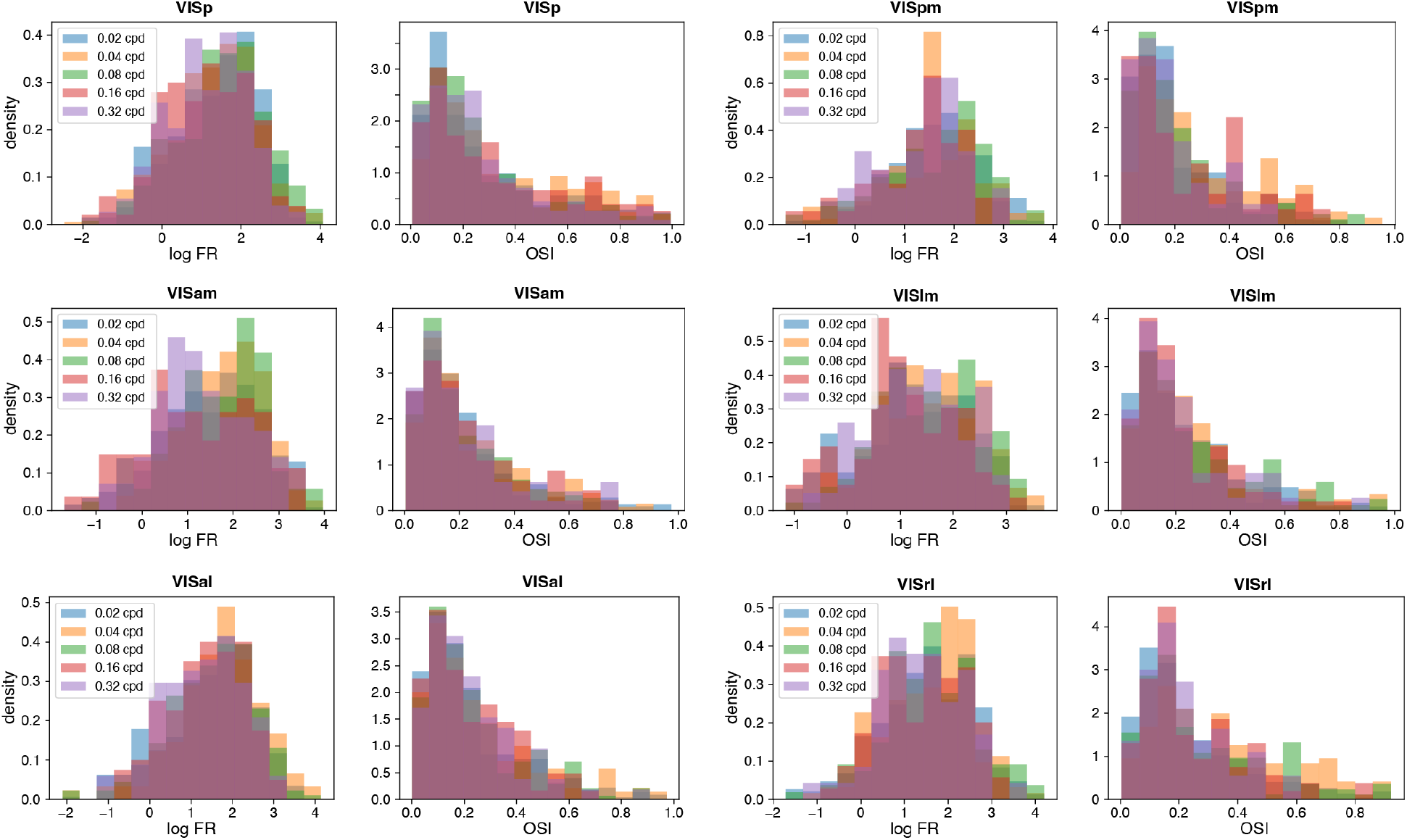
Analysis of spatial frequency preferences based on orientation selectivity and firing rates alone. A naive, histogram-based analysis of spatial frequency preferences based on traditional cell properties such as firing rate (FR) and orientation selectivity index (OSI) does not reveal clear differences or similarities between visual areas (compare with Figure 6a,b and Figures S2 and S13). This highlights the advantages of using data-driven, unsupervised approaches such as the encoding manifold method for understanding how functional properties organize across a large neuronal population.

**Fig. S17:**
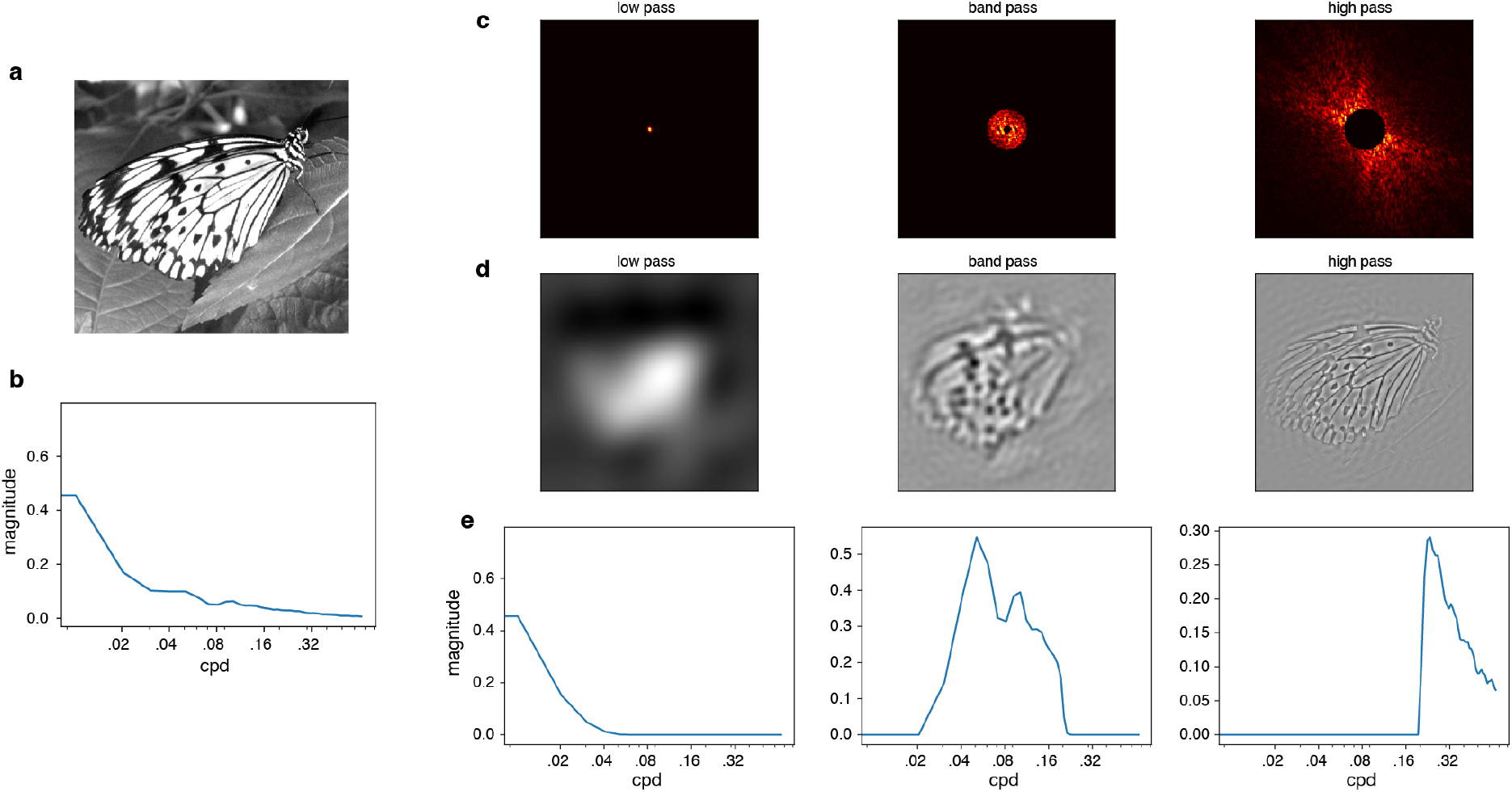
Filtering natural scenes. **a**, An example natural scene and **b**, its corresponding spatial frequency spectrum (azimuthal average), in cycles/degree (cpd) of visual angle. **c**, Images were filtered by cropping their spectra in Fourier domain (see Methods). **d** Resulting filtered images and **e** their corresponding spectra (azimuthal averages) after filtering was performed.

**Fig. S18:**
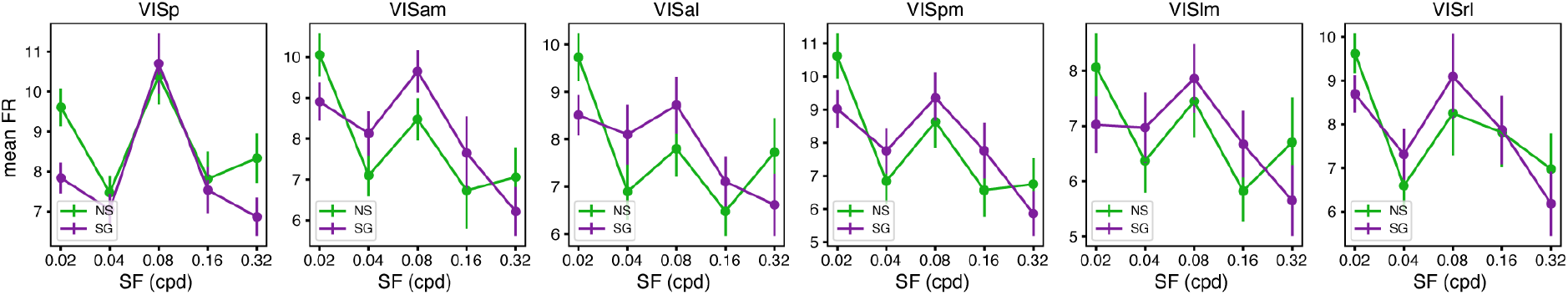
Mean firing rates to natural scenes and static gratings as a function of preferred spatial frequency. Mean *±* s.e.m. firing rates to natural scenes (NS; green) and static gratings (SG; magenta) are shown separately as a function of each neuron’s preferred spatial frequency, for all visual cortical areas. Neurons preferring the lowest (0.02 cpd) and highest (0.32 cpd) spatial frequencies tended to exhibit larger responses to natural scenes than to static gratings, whereas neurons preferring intermediate spatial frequencies generally responded more strongly to static gratings. These plots provide a complementary view of the U-shaped relationship between preferred spatial frequency and the log response ratio between NS and SG shown in Fig 7b in the main text. Note that these curves are computed from population mean firing rates and therefore are not mathematically equivalent to the analysis in Fig 7b, which reports the mean of per-neuron log response ratios.

## References

[1] E. Acar, D. M. Dunlavy, and T. G. Kolda. A scalable optimization approach for fitting canonical tensor decompositions. J Chemom, 25(2):67–86, 2011. doi: 10.1002/cem.1335.

[2] E. A. Allen and R. D. Freeman. Dynamic spatial processing originates in early visual pathways. Journal of Neuroscience, 26(45):11763–11774, 2006.

[3] Allen Institute. Allen Brain Observatory – Neuropixels Visual Coding. https://brainmapportal-live-4cc80a57cd6e400d854-f7fdcae.divio-media.net/filer_public/80/75/8075a100-ca64-429a-b39a-569121b612b2/neuropixels_visual_coding_-_white_paper_v10.pdf, 2019. Version 1.0 (October 1, 2019) Accessed: 02–Feb–2026].

[4] Allen Institute. Visual coding – neuropixels. https://portal.brain-map.org/circuits-behavior/visual-coding-neuropixels, 2024. Accessed: 02–Feb–2026.

[5] Allen Institute. Allen brain atlas software development kit (allensdk). https://allensdk.readthedocs.io/en/latest/visual_coding_neuropixels.html, 2024. Accessed: 02–Feb–2026.

[6] J.-M. Alonso, W. M. Usrey, and R. C. Reid. Precisely correlated firing in cells of the lateral geniculate nucleus. Nature, 383(6603):815–819, 1996.

[7] B. W. Bader and T. G. Kolda. Efficient MATLAB computations with sparse and factored tensors. SIAM J Sci Comput, 30(1):205–231, 2008.

[8] J. Bertram, L. Dyballa, T. A. Keller, S. Kinger, and S. W. Zucker. Manifolds and modules: How function develops in a neural foundation model. In Data on the Brain & Mind Workshop at the Neural Information Processing Systems Conference (NeurIPS), 2025.

[9] A. Blot, M. M. Roth, I. Gasler, M. Javadzadeh, F. Imhof, and S. B. Hofer. Visual intracortical and transthalamic pathways carry distinct information to cortical areas. Neuron, 109(12):1996–2008, 2021.

[10] F. Bolanños, J. G. Orlandi, R. Aoki, A. V. Jagadeesh, J. L. Gardner, and A. Benucci. Efficient coding of natural images in the mouse visual cortex. Nature Communications, 15(1):2466, 2024.

[11] G. Born, F. A. Schneider-Soupiadis, S. Erisken, A. Vaiceliunaite, C. L. Lao, M. H. Mobarhan, M. A. Spacek, G. T. Einevoll, and L. Busse. Corticothalamic feedback sculpts visual spatial integration in mouse thalamus. Nature Neuroscience, 24(12):1711–1720, 2021.

[12] Z. I. Botev, J. F. Grotowski, and D. P. Kroese. Kernel density estimation via diffusion. Ann Stat, 38 (5):2916–2957, 2010.

[13] C. E. Bredfeldt and D. Ringach. Dynamics of spatial frequency tuning in macaque v1. Journal of Neuroscience, 22(5):1976–1984, 2002.

[14] R. Bro and S. De Jong. A fast non-negativity-constrained least squares algorithm. J Chemom, 11(5): 393–401, 1997.

[15] S. Chung and L. Abbott. Neural population geometry: An approach for understanding biological and artificial neural networks. Curr Opin Neurobiol, 70:137–144, 2021.

[16] A. Cichocki, R. Zdunek, A. H. Phan, and S.-i. Amari. Nonnegative matrix and tensor factorizations: applications to exploratory multi-way data analysis and blind source separation. John Wiley & Sons, 2009.

[17] R. R. Coifman and S. Lafon. Diffusion maps. Appl Comput Harmon Anal, 21(1):5–30, 2006.

[18] R. R. Coifman, S. Lafon, A. B. Lee, M. Maggioni, B. Nadler, F. Warner, and S. W. Zucker. Geometric diffusions as a tool for harmonic analysis and structure definition of data: Diffusion maps. Proc Natl Acad Sci USA, 102(21):7426–7431, 2005.

[19] S. E. de Vries, J. A. Lecoq, M. A. Buice, P. A. Groblewski, G. K. Ocker, M. Oliver, D. Feng, N. Cain, P. Ledochowitsch, D. Millman, et al. A large-scale standardized physiological survey reveals functional organization of the mouse visual cortex. Nature Neuroscience, 23(1):138–151, 2020.

[20] S. E. de Vries, J. H. Siegle, and C. Koch. Sharing neurophysiology data from the allen brain observatory. eLife, 12:e85550, 2023.

[21] A. Dobbins, S. W. Zucker, and M. S. Cynader. Endstopped neurons in the visual cortex as a substrate for calculating curvature. Nature, 329(6138):438, 1987.

[22] L. Dyballa and S. W. Zucker. IAN: Iterated Adaptive Neighborhoods for manifold learning and dimensionality estimation. Neural Comput, 35, 2023.

[23] L. Dyballa, M. S. Hoseini, M. C. Dadarlat, S. W. Zucker, and M. P. Stryker. Flow stimuli reveal ecologically appropriate responses in mouse visual cortex. Proc Natl Acad Sci USA, 115(44):11304– 11309, 2018.

[24] L. Dyballa, A. M. Rudzite, M. S. Hoseini, M. Thapa, M. P. Stryker, G. D. Field, and S. W. Zucker. Population encoding of stimulus features along the visual hierarchy. Proceedings of the National Academy of Sciences, 121(4):e2317773121, 2024.

[25] E. Froudarakis, P. Berens, A. S. Ecker, R. J. Cotton, F. H. Sinz, D. Yatsenko, P. Saggau, M. Bethge, and A. S. Tolias. Population code in mouse V1 facilitates readout of natural scenes through increased sparseness. Nat Neurosci, 17(6):851, 2014.

[26] L. L. Glickfeld and S. R. Olsen. Higher-order areas of the mouse visual cortex. Annual Review of Vision Science, 3:251–273, 2017.

[27] Y. Han, J. M. Kebschull, R. A. Campbell, D. Cowan, F. Imhof, A. M. Zador, and T. D. Mrsic-Flogel. The logic of single-cell projections from visual cortex. Nature, 556(7699):51–56, 2018.

[28] C. R. Harris, K. J. Millman, S. J. Van Der Walt, R. Gommers, P. Virtanen, D. Cournapeau, et al. Array programming with NumPy. Nature, 585(7825):357–362, Sept. 2020. doi: 10.1038/s41586-020-2649-2. URL https://doi.org/10.1038/s41586-020-2649-2.

[29] J. A. Harris, S. Mihalas, K. E. Hirokawa, J. D. Whitesell, H. Choi, A. Bernard, P. Bohn, S. Caldejon, L. Casal, A. Cho, et al. Hierarchical organization of cortical and thalamic connectivity. Nature, 575 (7781):195–202, 2019.

[30] J. Hegdé. Time course of visual perception: coarse-to-fine processing and beyond. Progress in neurobiology, 84(4):405–439, 2008.

[31] J. L. Hoy, I. Yavorska, M. Wehr, and C. M. Niell. Vision drives accurate approach behavior during prey capture in laboratory mice. Curr Biol, 26(22):3046–3052, 2016.

[32] S. Katzner, G. Born, and L. Busse. V1 microcircuits underlying mouse visual behavior. Current opinion in neurobiology, 58:191–198, 2019.

[33] A. J. Keller, M. M. Roth, and M. Scanziani. Feedback generates a second receptive field in neurons of the visual cortex. Nature, 582(7813):545–549, 2020.

[34] A. M. Kerlin, M. L. Andermann, V. K. Berezovskii, and R. C. Reid. Broadly tuned response properties of diverse inhibitory neuron subtypes in mouse visual cortex. Neuron, 67(5):858–871, 2010.

[35] E. J. Kim, A. L. Juavinett, E. M. Kyubwa, M. W. Jacobs, and E. M. Callaway. Three types of cortical layer 5 neurons that differ in brain-wide connectivity and function. Neuron, 88(6):1253–1267, 2015.

[36] M.-H. Kim, P. Znamenskiy, M. F. Iacaruso, and T. D. Mrsic-Flogel. Segregated subnetworks of intracortical projection neurons in primary visual cortex. Neuron, 100(6):1313–1321, 2018.

[37] S. Kinger, L. Dyballa, and S. W. Zucker. Bracketing uncertainty in clustering under the manifold hypothesis. arXiv preprint, 2026.

[38] L. Kirchberger, S. Mukherjee, U. H. Schnabel, E. H. van Beest, A. Barsegyan, C. N. Levelt, J. A. Heimel, J. A. Lorteije, C. van der Togt, M. W. Self, et al. The essential role of recurrent processing for figure-ground perception in mice. Science Advances, 7(27):eabe1833, 2021.

[39] M. A. Kirchgessner, A. D. Franklin, and E. M. Callaway. Distinct “driving” versus “modulatory” influences of different visual corticothalamic pathways. Current Biology, 31(23):5121–5137, 2021.

[40] T. G. Kolda. Multilinear operators for higher-order decompositions. Technical Report SAND2006-2081, Sandia National Laboratories, April 2006. URL http://www.osti.gov/scitech/biblio/923081.

[41] C. Kong, Y. Wang, and G. Xiao. Neuron populations across layer 2-6 in the mouse visual cortex exhibit different coding abilities in the awake mice. Frontiers in Cellular Neuroscience, 17:1238777, 2023.

[42] D. D. Lee and H. S. Seung. Learning the parts of objects by non-negative matrix factorization. Nature, 401(6755):788–791, 1999.

[43] F. J. Luongo, L. Liu, C. L. A. Ho, J. K. Hesse, J. B. Wekselblatt, F. F. Lanfranchi, D. Huber, and D. Y. Tsao. Mice and primates use distinct strategies for visual segmentation. eLife, 12:e74394, 2023.

[44] W.-P. Ma, B.-H. Liu, Y.-T. Li, Z. J. Huang, L. I. Zhang, and H. W. Tao. Visual representations by cortical somatostatin inhibitory neurons–selective but with weak and delayed responses. J. Neurosci., 30(43):14371–14379, Oct. 2010.

[45] J. Manley, S. Lu, K. Barber, J. Demas, H. Kim, D. Meyer, F. M. Traub, and A. Vaziri. Simultaneous, cortex-wide dynamics of up to 1 million neurons reveal unbounded scaling of dimensionality with neuron number. Neuron, 2024.

[46] D. Marr and T. Poggio. A computational theory of human stereo vision. Proceedings of the Royal Society B: Biological Sciences, 204(1156):301–328, 1979.

[47] J. H. Marshel, M. E. Garrett, I. Nauhaus, and E. M. Callaway. Functional specialization of seven mouse visual cortical areas. Neuron, 72(6):1040–1054, 2011.

[48] D. Martin, C. Fowlkes, D. Tal, and J. Malik. A database of human segmented natural images and its application to evaluating segmentation algorithms and measuring ecological statistics. In Proceedings of the 8th IEEE International Conference on Computer Vision (ICCV 2001), volume 2, pages 416–423. IEEE, 2001.

[49] J. A. Mazer, W. E. Vinje, J. McDermott, P. H. Schiller, and J. L. Gallant. Spatial frequency and orientation tuning dynamics in area v1. Proceedings of the National Academy of Sciences, 99(3): 1645–1650, 2002.

[50] L. McInnes, J. Healy, N. Saul, and L. Großberger. UMAP: Uniform manifold approximation and projection. Journal of Open Source Software, 3(29):861, 2018. doi: 10.21105/joss.00861.

[51] J. S. Montijn, P. M. Goltstein, and C. M. Pennartz. Mouse v1 population correlates of visual detection rely on heterogeneity within neuronal response patterns. Elife, 4:e10163, 2015.

[52] P. Murphy and A. Sillito. Corticofugal feedback influences the generation of length tuning in the visual pathway. Nature, 329(6141):727–729, 1987.

[53] C. M. Niell and M. Scanziani. How cortical circuits implement cortical computations: mouse visual cortex as a model. Annual Review of Neuroscience, 44:517–546, 2021.

[54] C. M. Niell and M. P. Stryker. Highly selective receptive fields in mouse visual cortex. J Neurosci, 28 (30):7520–7536, 2008.

[55] T. Odland. KDEpy: Kernel Density Estimation in Python, Dec. 2018. v0.9.10. Available at 10.5281/zenodo.2392268.

[56] A. Oliva and A. Torralba. Modeling the shape of the scene: A holistic representation of the spatial envelope. International Journal of Computer Vision, 42:145–175, 2001.

[57] A. Olmos and F. A. Kingdom. A biologically inspired algorithm for the recovery of shading and reflectance images. Perception, 33(12):1463–1473, 2004.

[58] R. V. Rikhye and M. Sur. Spatial correlations in natural scenes modulate response reliability in mouse visual cortex. J Neurosci, 35(43):14661–14680, 2015.

[59] J. H. Siegle, X. Jia, S. Durand, S. Gale, C. Bennett, N. Graddis, G. Heller, T. K. Ramirez, H. Choi, J. A. Luviano, et al. Survey of spiking in the mouse visual system reveals functional hierarchy. Nature, pages 1–7, 2021.

[60] A. M. Sillito, H. E. Jones, G. L. Gerstein, and D. C. West. Feature-linked synchronization of thalamic relay cell firing induced by feedback from the visual cortex. Nature, 369(6480):479–482, 1994.

[61] R. Skyberg, S. Tanabe, H. Chen, and J. Cang. Coarse-to-fine processing drives the efficient coding of natural scenes in mouse visual cortex. Cell reports, 38(13), 2022.

[62] R. J. Skyberg and C. M. Niell. Natural visual behavior and active sensing in the mouse. Current Opinion in Neurobiology, 86:102882, 2024.

[63] I. T. Smith, L. B. Townsend, R. Huh, H. Zhu, and S. L. Smith. Stream-dependent development of higher visual cortical areas. Nature Neuroscience, 20(2):200–208, 2017.

[64] C. Stringer, M. Michaelos, D. Tsyboulski, S. E. Lindo, and M. Pachitariu. High-precision coding in visual cortex. Cell, 184(10):2767–2778, 2021.

[65] E. Switkes, M. J. Mayer, and J. A. Sloan. Spatial frequency analysis of the visual environment: Anisotropy and the carpentered environment hypothesis. Vision Research, 18(10):1393–1399, 1978.

[66] L. Van der Maaten and G. Hinton. Visualizing data using t-sne. J Mach Learn Res, 9(11), 2008.

[67] J. H. Van Hateren and A. van der Schaaf. Independent component filters of natural images compared with simple cells in primary visual cortex. Proceedings of the Royal Society of London. Series B: Biological Sciences, 265(1394):359–366, 1998.

[68] M. Vélez-Fort, C. V. Rousseau, C. J. Niedworok, I. R. Wickersham, E. A. Rancz, A. P. Brown, M. Strom, and T. W. Margrie. The stimulus selectivity and connectivity of layer six principal cells reveals cortical microcircuits underlying visual processing. Neuron, 83(6):1431–1443, 2014.

[69] S. Vreysen, B. Zhang, Y. M. Chino, L. Arckens, and G. Van den Bergh. Dynamics of spatial frequency tuning in mouse visual cortex. Journal of neurophysiology, 107(11):2937–2949, 2012.

[70] Q. Wang, O. Sporns, and A. Burkhalter. Network analysis of corticocortical connections reveals ventral and dorsal processing streams in mouse visual cortex. Journal of Neuroscience, 32(13):4386–4399, 2012.

[71] F. A. Wichmann, J. Drewes, P. Rosas, and K. R. Gegenfurtner. Animal detection in natural scenes: Critical features revisited. Journal of Vision, 10(4):6–6, 2010.

[72] A. H. Williams, T. H. Kim, F. Wang, S. Vyas, S. I. Ryu, K. V. Shenoy, M. Schnitzer, T. G. Kolda, and S. Ganguli. Unsupervised discovery of demixed, low-dimensional neural dynamics across multiple timescales through tensor component analysis. Neuron, 98(6):1099–1115, 2018.

[73] Y. Yu, J. N. Stirman, C. R. Dorsett, and S. L. Smith. Selective representations of texture and motion in mouse higher visual areas. Current Biology, 32(13):2810–2820, 2022.

[74] Y. Yu, J. N. Stirman, C. R. Dorsett, and S. L. Smith. Visual information is broadcast among cortical areas in discrete channels. bioRxiv, pages 2023–12, 2023.

[75] H. Zhou, R. J. Schafer, and R. Desimone. Pulvinar-cortex interactions in vision and attention. Neuron, 89(1):209–220, 2016.

